# An intermembrane space protein facilitates completion of mitochondrial division in yeast

**DOI:** 10.1101/2023.03.31.535139

**Authors:** Olivia M. Connor, Srujan K. Matta, Jonathan R. Friedman

## Abstract

Mitochondria are highly dynamic double membrane-bound organelles that maintain their shape in part through fission and fusion. Mitochondrial fission is performed by the dynamin-related protein Dnm1 (Drp1 in humans), a large GTPase that constricts and divides the mitochondria in a GTP hydrolysis-dependent manner. However, it is unclear whether factors inside mitochondria help coordinate the process and if Dnm1/Drp1 activity alone is sufficient to complete fission of both mitochondrial membranes. Here, we identify an intermembrane space protein required for mitochondrial fission in yeast, which we propose to name Mdi1. Loss of Mdi1 leads to hyper-fused mitochondria networks due to defects in mitochondrial fission, but not lack of Dnm1 recruitment to mitochondria. Mdi1 plays a conserved role in fungal species and its homologs contain a putative amphipathic α-helix, mutations in which disrupt mitochondrial morphology. One model to explain these findings is that Mdi1 associates with and distorts the mitochondrial inner membrane to enable Dnm1 to robustly complete fission. Our work reveals that Dnm1 cannot efficiently divide mitochondria without the coordinated function of a protein that resides inside mitochondria.

## Introduction

Mitochondria are multifunctional organelles distributed throughout cells as part of a dynamic network of tubules (Friedman and Nunnari, 2014). Mitochondrial subcellular localization is in part controlled by the ability of individual mitochondria to fuse together or undergo fission. These opposing conserved processes are in balance at steady-state, and disruption of either fusion or fission leads to altered distribution of mitochondria, disrupted mitochondrial DNA (mtDNA) maintenance, and respiratory dysfunction (Quintana-Cabrera and Scorrano, 2023). Mitochondrial division is also important for organelle quality control and cellular homeostasis as it is associated with the removal of damaged portions of the mitochondrial network during mitophagy (Pickles et al., 2018).

Mitochondria are double membrane-bound and the fission/fusion dynamics of the organelle necessitates remodeling of both the mitochondrial inner membrane (IMM) and the mitochondrial outer membrane (OMM). In accordance with the fundamental importance of mitochondrial dynamics, the processes of fission and fusion occur in unicellular eukaryotes and metazoan species and the core machineries that perform these functions are conserved (Gao and Hu, 2021; Kraus et al., 2021). Both fission and fusion are mediated by members of a family of large GTPases, the dynamin-related proteins (DRPs). In the case of mitochondrial fusion, this requires the coordinated sequential action of fusion DRPs on the OMM and IMM (Gao and Hu, 2021). During mitochondrial division, the fission DRP (Dnm1 in yeast, Drp1 in humans) is recruited to the OMM by receptor proteins, assembles in an oligomeric structure that circumscribes mitochondria, and constricts and mediates fission of the membranes via a confirmational change driven by GTP hydrolysis (Kraus et al., 2021).

Several factors contribute to the spatial regulation of mitochondrial division. Inter-organelle contacts, primarily between the ER and mitochondria, mark sites of membrane pre-constriction that precedes mitochondrial fission in both yeast and humans (Friedman et al., 2011; Guo et al., 2018; Kleele et al., 2021). In human cells, the ER membrane phospholipid modifying enzyme ABHD16A may contribute to mitochondrial constrictions by acting *in trans* on the mitochondrial surface (Nguyen and Voeltz, 2022), though the ER may also promote mitochondrial constrictions via INF2-dependent polymerization of actin at the membrane contact site (MCS) (Chakrabarti et al., 2018; Korobova et al., 2013; Manor et al., 2015). In addition to external determinants, fission of the organelle must also be coordinated with factors inside mitochondria. Sites of replication of mtDNA, which occurs in the matrix compartment interior to the IMM, are spatially linked to sites of mitochondrial division in human cells (Lewis et al., 2016). However, despite the relationship between both inter-organelle contacts and replication of mtDNA to mitochondrial division sites, it is still unknown how the processes are spatially coordinated across the two mitochondrial membrane bilayers and whether factors inside mitochondria play a direct role in facilitating mitochondrial division. Additionally, it has remained an open question whether the activity of the Dnm1/Drp1 division GTPase is sufficient to fully drive fission of both mitochondrial membranes, whether a separate IMM fission machinery exists, or if other external factors assist in completing membrane scission (Anand et al., 2014; Chakrabarti et al., 2018; Fonseca et al., 2019; Kamerkar et al., 2018; Klecker et al., 2015; Lee et al., 2016). For example, the yeast IMM protein Mdm33 has been implicated as an auxiliary mediator of mitochondrial division, though the absence of Mdm33 does not lead to mitochondrial morphology defects that phenocopy loss of other known mitochondrial fission proteins (Klecker et al., 2015; Messerschmitt et al., 2003).

In recent years, a number of in-depth proteomics analyses in yeast and metazoans have generated a more complete inventory of the mitochondrial proteome (Hung et al., 2014; Morgenstern et al., 2017; Rath et al., 2021; Rensvold et al., 2022; Rhee et al., 2013; Stefely et al., 2016; Vogtle et al., 2017). These studies and others have identified smaller proteins (often defined as less than 100 amino acids) called micropeptides or small Open Reading Frames (smORFs), that localize inside mitochondria (Tharakan and Sawa, 2021; Zheng and Xiang, 2022). In metazoans, these internal mitochondrial micropeptides have been frequently implicated in critical functions such as the assembly of respiratory complexes (Bosch et al., 2022; Chu et al., 2019; Rathore et al., 2018; Zhang et al., 2020). In yeast, recent work has experimentally validated the presence of many such micropeptides, begging the question of their functional roles (Morgenstern et al., 2017).

Here, we characterize one such micropeptide in budding yeast, a 73 amino acid protein named Mco8 (Mitochondria Class One 8 kDa; named for its molecular weight and its high confidence localization to mitochondria), which was previously validated as an intermembrane space (IMS) protein (Morgenstern et al., 2017). We show that Mco8 can be spatially linked to sites of mitochondrial fission and its loss leads to a disruption of the process, prompting us to propose the name <u>M</u>itochondrial <u>D</u>ivision IMS 1 (Mdi1). We find loss of Mdi1 likely leads to hyper-fused mitochondria due to an inability of Dnm1 to complete scission of the organelle. While Mdi1 is absent in metazoans, it is present throughout fungal species, and here we show that it plays a conserved role in fission yeast. Mdi1 is characterized by a structurally conserved putative amphipathic α-helix, and we find mutations in this motif prevent its function in maintaining normal mitochondrial morphology. These data are consistent with a model that Mdi1 locally binds and/or distorts mitochondrial membranes from the inside of the organelle, enabling Dnm1 to complete mitochondrial fission. Thus, we have identified the first mitochondrial IMS-localized factor directly involved in the process of mitochondrial division.

## Results

### Mitochondrial division is perturbed in the absence of Mdi1

To explore a functional role for the IMS-targeted micropeptide Mdi1, we examined mitochondrial morphology in Δ*mdi1* yeast cells using fluorescence microscopy. In contrast to wild type cells, which exhibit tubular mitochondrial morphology, mitochondria in Δ*mdi1* cells had an aberrant net-like appearance in nearly all cells, reminiscent of cells with hyper-fused mitochondrial networks (Fig. 1A). We thus compared mitochondrial morphology in Δ*mdi1* cells to those of cells deficient for the mitochondrial division DRP, Dnm1 (Δ*dnm1*), or its OMM receptor, Fis1 (*Δfis1*). Mitochondrial morphology appeared similarly net-like in Δ*mdi1* cells as in Δ*dnm1* or Δ*fis1* cells (Fig. 1A-1B).

**Figure 1.**
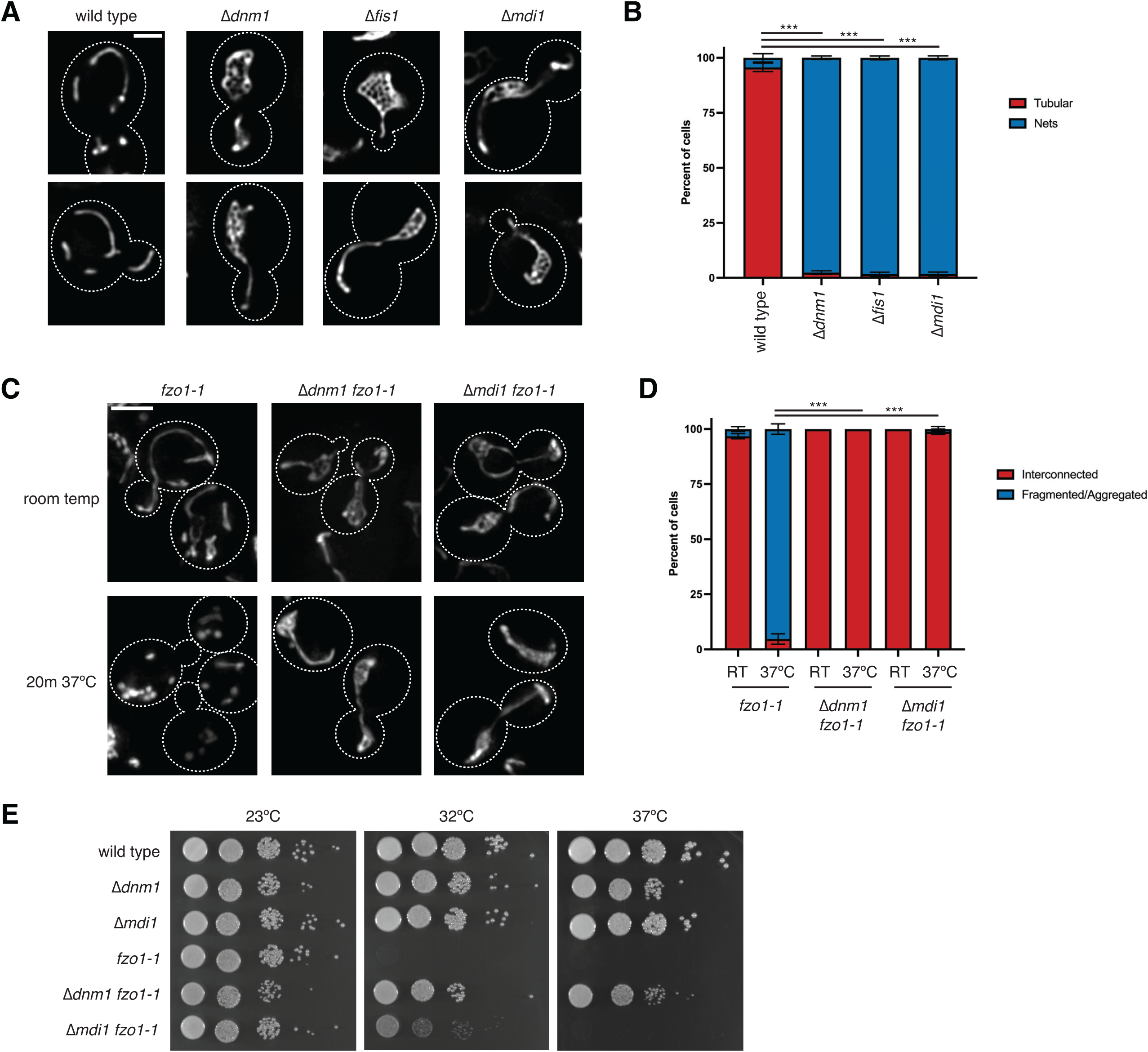
Mitochondrial division is perturbed in the absence of Mdi1. **(A)** Representative single plane deconvolved epifluorescence microscopy images of the indicated budding yeast strains expressing the mitochondrial matrix marker mito-dsRed. **(B)** A graph of the categorization of mitochondrial morphology from cells as in (A). Data shown represent at least 75 cells per strain in each of three independent experiments and bars indicate S.E.M. Asterisks (***p<0.001) represent unpaired two-tailed *t* tests. **(C)** Maximum intensity projections of epifluorescence microscopy images of the indicated strains expressing mito-dsRed that were grown to log phase at room temperature and imaged before (top) and after (bottom) growth at 37°C for 20 minutes. **(D)** A graph depicting categorization of mitochondrial morphology as in (C). Data shown represent at least 70 cells per strain in each of three independent experiments and bars indicate S.E.M. Asterisks (***p<0.001) represent unpaired two-tailed *t* tests. **(E)** Serial dilutions of the indicated yeast cells plated on media containing the non-fermentable carbon source ethanol/glycerol (YPEG) and grown at the indicated temperatures. Cell boundaries are indicated with dotted lines. Scale bars = 2 µm (A), 3 µm (C). See also Figure S1.

Tubular mitochondrial networks are formed by a balance between opposing fission and fusion processes (Bleazard et al., 1999; Nunnari et al., 1997; Sesaki and Jensen, 1999), and we reasoned that the net-like morphology caused by loss of Mdi1 could be due either to a deficiency in mitochondrial fission or to excessive mitochondrial fusion. To address this, we utilized a temperature-sensitive allele of the gene encoding the mitochondrial OMM fusion DRP Fzo1 (*fzo1-1*; (Hermann et al., 1998)). Consistent with published observations, mitochondria in *fzo1-1* cells maintained wild type tubular appearance when grown at room temperature (Fig. 1C). However, growth at a non-permissive temperature (37°C) causes deficiency in Fzo1-dependent mitochondrial fusion, and within 20 minutes, mitochondria appeared fragmented and/or aggregated in the majority of cells (Fig. 1C-1D). We then asked how loss of Dnm1 or Mdi1 impacted mitochondrial morphology in the *fzo1-1* yeast strain. As expected, mitochondria in Δ*dnm1 fzo1-1* cells did not appear fragmented at an elevated temperature and instead, the mitochondria appeared tubular and/or net-like in all cells (Fig. 1C-1D; (Mozdy et al., 2000)). Remarkably, Δ*mdi1 fzo1-1* cells similarly maintained interconnected mitochondria in nearly all cells, similar to loss of Dnm1 (Fig. 1C-1D). These data suggest that Mdi1 plays a positive role in mitochondrial division.

Mitochondrial fission/fusion dynamics are required for the maintenance of mtDNA, and excessive mitochondrial fragmentation due to a lack of fusion leads to mtDNA loss and therefore an inability for yeast cells to grow on media requiring mitochondrial respiration (Hermann et al., 1998). However, simultaneous loss of both division and fusion machinery prevents mitochondrial fragmentation and allows for genome maintenance, albeit with an increased rate of mtDNA mutation and loss (Bleazard et al., 1999; Mozdy et al., 2000; Osman et al., 2015; Tieu and Nunnari, 2000). Thus, while *fzo1-1* yeast cells can grow on media containing a carbon source that requires mitochondrial respiration (ethanol/glycerol) at the permissive temperature (23°C), they are inviable at elevated temperatures (Fig. 1E; (Hermann et al., 1998)). However, in the absence of fission machinery such as Dnm1, mitochondria remain interconnected even when Fzo1 is inactivated, enabling mtDNA maintenance and growth on ethanol/glycerol media (Fig. 1E). We therefore asked whether Δ*mdi1 fzo1-1* cells were viable on respiration-requiring media grown at a non-permissive temperature. We found that while the cells remained inviable at 37°C, loss of Mdi1 promoted growth to *fzo1-1* cells at an intermediate temperature, 32°C (Fig. 1E). These data suggest that in cells lacking Mdi1, mitochondrial fission is deficient, though may not be completely absent.

Loss of mitochondrial membrane potential has previously been established to induce rapid Dnm1-dependent mitochondrial fission in yeast (Hughes et al., 2016; Klecker et al., 2015). As a complementary approach, we asked whether loss of Mdi1 prevented mitochondrial fragmentation induced by treatment with the uncoupler CCCP, which would suggest a deficiency in mitochondrial division. We treated cells with CCCP (25µM, 45 minutes), and as expected, the mitochondria in the majority of wild type cells appeared fragmented (Fig. S1A-S1B). In contrast, loss of Dnm1 prevented mitochondrial fragmentation in most cells and the networks remained interconnected. Notably, mitochondria in Δ*dnm1* cells treated with CCCP frequently appeared hyper-constricted in a beads-on-a-string appearance, indicative of Dnm1-independent mitochondrial constriction (Legesse-Miller et al., 2003) (Fig. S1A-S1B, see arrows). As in Δ*dnm1* cells, loss of Mdi1 appeared to largely prevent mitochondrial fragmentation, though not hyper-constriction, upon CCCP treatment (Fig. S1A-S1B, see arrows). Altogether, these data indicate that loss of Mdi1 leads to hyper-fused mitochondria due to a deficiency in mitochondrial fission.

### Mdi1 is a soluble intermembrane space protein

Previous work characterizing Mdi1 included analysis of its sub-mitochondrial localization via a biochemical protease protection assay that relied on a C-terminal Protein A fusion to enable its detection, the results of which indicated that the protein resides in the IMS (Morgenstern et al., 2017). As Mdi1 is not suggested to have transmembrane domains according to hydrophobicity/transmembrane prediction programs such as TMHMM and Phobius (Kall et al., 2004; Krogh et al., 2001), Mdi1 likely influences Dnm1-dependent mitochondrial fission from inside the organelle. To our knowledge, depletion of no other internal mitochondrial factor in yeast leads to a net-like mitochondrial morphology that phenocopies loss of Dnm1, which motivated us to confirm its localization.

In our hands, several different chromosomally integrated C-terminal fusions to Mdi1 led to mitochondrial morphology defects similar to Δ*mdi1* cells, indicating that these tags interfere with its function. Therefore, to confirm the sub-mitochondrial localization of Mdi1, we generated an in-frame 2xFLAG tag fusion (Mdi1*-2xFLAG) in a putative unstructured region of the protein based on AlphaFold 2 predictions (Fig. S2A; (Jumper et al., 2021; Varadi et al., 2022)). This fusion protein, when integrated at the *ura3* locus and expressed using the native *MDI1* promoter, was capable of fully rescuing the mitochondrial morphology defect of Δ*mdi1* cells (Fig. S2B).

As the AlphaFold 2 prediction suggests Mdi1 contains two lengthy alpha-helical regions separated by a short linker (Fig. S2A), we considered that the protein may be an integral membrane protein. However, treatment of mitochondria isolated from Mdi1*-2xFLAG expressing cells with an alkaline solution of sodium carbonate indicates that Mdi1 could readily be extracted from membranes, suggesting it is a soluble protein (Fig. S2C). Further, consistent with published work, a protease protection analysis with tagged functional Mdi1 suggests that it is protected from proteolytic cleavage unless the OMM is disrupted, similar to the soluble IMS protein Mia40 (Fig. S2D; (Chacinska et al., 2004)). Thus, these data indicate that Mdi1 is a soluble, IMS-localized protein.

### Mdi1 localizes at discrete focal structures that can be spatially linked to sites of mitochondrial division

Given the genetic involvement of Mdi1 in mitochondrial fission, we next wanted to ascertain Mdi1 sub-localization within mitochondria and its spatial relationship to mitochondrial division sites. Mdi1 was nearly impossible to detect when we tagged it with a single fluorescent protein. We circumvented this issue by C-terminally tagging Mdi1 with a 7x tandem GFP11 tag, which we could then visualize by artificially targeting the complementary GFP1-10 to the IMS via the N-terminal sequence of Cyb2 (Fig. 2A) (Beasley et al., 1993; Kamiyama et al., 2016). We observed Mdi1-GFP11_x7_ localized to discrete focal structures relative to the matrix marker mito-dsRed (Fig. 2B, arrows). However, the tag renders Mdi1 non-functional for mitochondrial division as expression of the fusion could not rescue mitochondrial morphology of Δ*mdi1* cells (Fig. 2B-2C). Despite this, the tagged protein does not interfere with mitochondrial division in the presence of a wild type copy of the *MDI1* gene, suggesting the tagged form of the protein may reflect the normal localization of the protein (Fig. 2B-2C). In cells expressing both wild type Mdi1 and Mdi1-GFP11_x7_, each expressed using the native *MDI1* promoter, the tagged protein again localized in discrete focal structures that were distributed throughout the tubular mitochondrial network (Fig. 2B, arrows).

**Figure 2.**
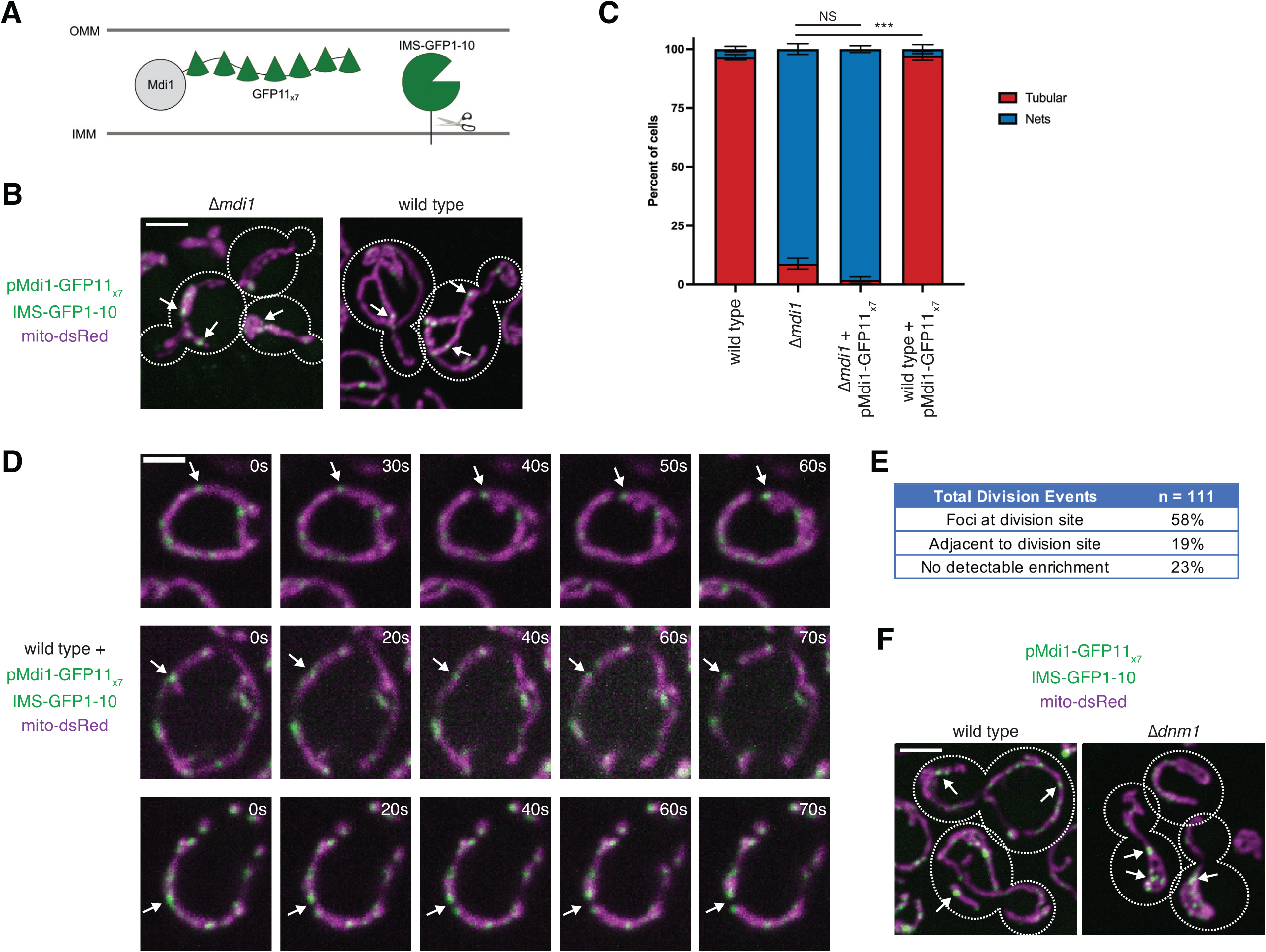
Mdi1 localizes at discrete focal structures that can be spatially linked to sites of mitochondrial division. **(A)** A schematic depicting the tandem tag split GFP approach utilized to visualize Mdi1. **(B)** Maximum intensity projections of confocal fluorescence microscopy images of Δ*mdi1* (left) or wild type (right) cells co-expressing IMS-targeted GFP1-10 and chromosomally integrated Mdi1-GFP11_x7_ driven by its native promoter (green) and mito-dsRed (magenta). Arrows mark focal accumulations of Mdi1-GFP11_x7_. **(C)** A graph depicting categorization of mitochondrial morphology from the indicated strains as in (B). Data shown represent at least 75 cells per strain in each of three independent experiments and bars indicate S.E.M. Asterisks (***p<0.001) represent unpaired two-tailed *t* test. NS indicates not statistically significant. **(D)** Timelapse microscopy images of wild type cells expressing Mdi1-GFP11_x7_ as in (B). Images shown are a maximum intensity projection of three confocal images with 0.4 µm z-steps captured at the indicated time intervals. Arrows mark sites of Mdi1-GFP11_x7_-marked mitochondrial division. **(E)** Categorization of the frequency that mitochondrial division events captured as in (D) were spatially linked to Mdi1-GFP11_x7_ foci. **(F)** Images as in (B) of wild type (left) or Δ*dnm1* (right) cells. Cell boundaries are indicated with dotted lines. Scale bars = 2 µm (D), 3 µm (B, F). See also Figure S2.

To examine the dynamics of Mdi1-GFP11_x7_ foci in cells co-expressing wild type Mdi1, we performed time-lapse confocal microscopy. While Mdi1-GFP11_x7_ is susceptible to photobleaching, we could regularly observe examples of discrete Mdi1 foci marking sites of mitochondrial division (Fig. 2D-2E, arrows; 77% of mitochondrial division events had Mdi1 spatially linked at or adjacent to the division site). With the caveat that the tagged form may not accurately reflect the behavior of the endogenous protein, these localization data in combination with our data indicating Mdi1 is required for efficient mitochondrial fission suggests that Mdi1 locally influences the division process.

We next asked how loss of Dnm1 impacted Mdi1 localization within mitochondria. We again exogenously expressed native-promoter driven Mdi1-GFP11_x7_ in either *MDI1 DNM1* cells or in a *MDI1* Δ*dnm1* strain background. In the absence of Dnm1, even though the mitochondria appeared net-like, Mdi1 retained its focal distribution within the mitochondrial network (Fig. 2F, arrows). Thus, the propensity of Mdi1-GFP11_x7_ to concentrate at mitochondrial division-linked structures appears to be Dnm1-independent.

### Dnm1 fails to complete mitochondrial division in the absence of Mdi1

Mitochondrial division requires the coordinated scission of both the OMM and the IMM of the organelle. However, several groups have reported evidence of the IMM constricting upstream and independently of Dnm1/Drp1 during mitochondria division in yeast, worms, mice, and mammalian cells (Chakrabarti et al., 2018; Cho et al., 2017; Friedman et al., 2011; Ishihara et al., 2009; Labrousse et al., 1999; Legesse-Miller et al., 2003). Indeed, we observed apparent mitochondrial constrictions during CCCP-induced mitochondrial division, even in the absence of Dnm1 (Fig. S1A, Fig. 3A). Given Mdi1 localization to the IMS and its ability to form discrete Dnm1-independent assemblies, we hypothesized that the protein could be responsible for constriction of the organelle upstream of Dnm1 recruitment. However, as in Δ*dnm1* cells, mitochondria in Δ*mdi1* cells also appear constricted after treatment with CCCP (Fig. S1A, Fig. 3A, arrows). We considered the possibility that in the absence of Mdi1, Dnm1 is sufficient to cause mitochondria constrictions, and therefore examined mitochondrial morphology in *Δmdi1* Δ*dnm1* cells. However, even in the absence of both Dnm1 and Mdi1, mitochondria in CCCP-treated cells appear hyper-constricted (Fig. 3A, arrows). Together, these data indicate that Dnm1-independent mitochondrial constrictions do not require Mdi1 and suggest an alternative role for the protein.

**Figure 3.**
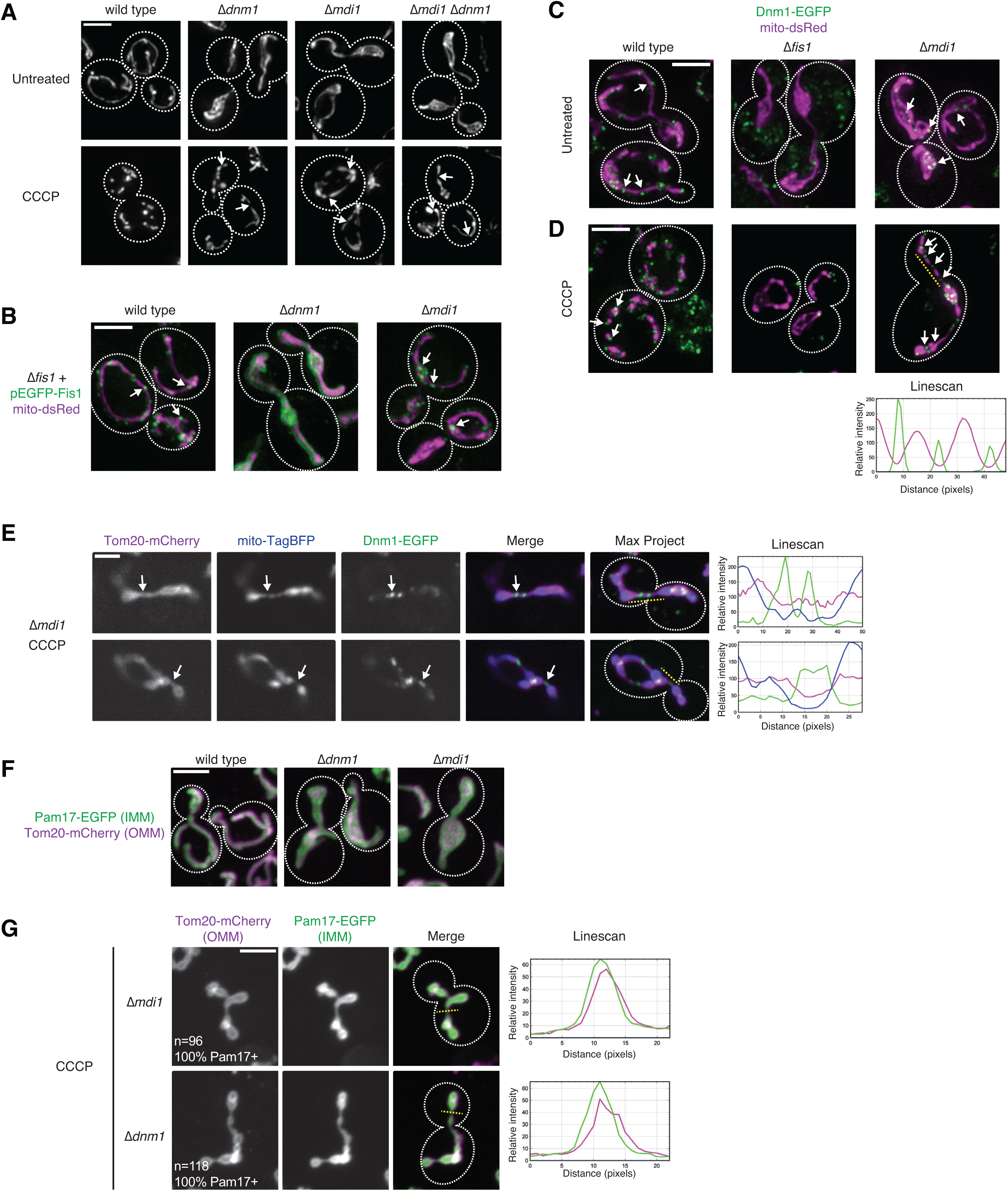
Dnm1 fails to complete mitochondrial division in the absence of Mdi1. **(A)** Maximum intensity projections of deconvolved epifluorescence microscopy images of the indicated yeast strains grown to log phase and either directly imaged (top) or treated for 40 minutes with 25 µM CCCP (bottom) prior to imaging. Arrows mark sites of mitochondrial constriction. **(B)** Maximum intensity projections of deconvolved epifluorescence microscopy images of the indicated yeast strains expressing chromosomally integrated EGFP-Fis1 (green) and mito-DsRed (magenta). Arrows mark sites of focal accumulation of Fis1. **(C)** As in (B) for cells expressing an endogenous chromosomal Dnm1-EGFP tag (green). Arrows mark sites of Dnm1 association with mitochondria. **(D)** As in (C) for cells treated for 30 minutes with 25 µM CCCP prior to imaging. Dashed yellow line corresponds to fluorescence intensity linescan shown below. **(E)** Single plane (left) and maximum intensity projection (right) confocal fluorescence microscopy images are shown of Δ*mdi1* cells expressing Tom20-mCherry (OMM, magenta), mito-TagBFP (matrix, blue), and Dnm1-EGFP (green). Arrows mark sites of Dnm1-marked hyper-constriction where the OMM label appears continuous. Dashed yellow lines correspond to the fluorescence intensity linescans shown at right. **(F)** Maximum intensity projections of confocal microscopy images of the indicated yeast strains expressing chromosomally tagged Tom20-mCherry (OMM, magenta) and Pam17-EGFP (IMM, green). **(G)** As in (F) for the indicated yeast strains treated for 45 minutes with 25 µM CCCP. Dashed yellow lines correspond to the fluorescence intensity linescans shown at right. Quantification shown in figure represents the percentage of Tom20-positive mitochondrial hyper-constrictions that appeared positive for Pam17-EGFP. Cell boundaries are indicated with dotted lines. Scale bars = 2 µm (E), 3 µm (A-D, F-G).

To gain further insight into the potential role of Mdi1 in mitochondrial division, we examined how its loss impacted the known mitochondrial fission machinery. We expressed chromosomally integrated EGFP-tagged Fis1, the OMM receptor for Dnm1, under the control of its native promoter in Δ*fis1* cells. In cells co-expressing the mitochondrial matrix marker mito-DsRed, Fis1 appeared distributed throughout the mitochondria but also enriched in discrete foci (Fig. 3B, arrows). Loss of Dnm1 caused Fis1 to redistribute and more uniformly decorate mitochondria, which is consistent with the notion that Fis1 focal localization is promoted by Dnm1 recruitment to the OMM (Fig. 3B). However, in contrast, GFP-Fis1 remained focal along the hyper-fused mitochondria in Δ*mdi1* cells, suggesting Dnm1 recruitment may not be affected (Fig. 3B, arrows).

We next examined Dnm1 localization with an endogenous chromosomal C-terminal EGFP fusion in cells co-expressing mito-dsRed. In wild type cells, Dnm1 appeared in focal structures that were predominantly associated with mitochondria (Fig. 3C, arrows). As expected, in Δ*fis1* cells, Dnm1 did not localize to mitochondria and instead appeared in discrete cytosolic foci (Fig. 3C) (Mozdy et al., 2000). In contrast, in Δ*mdi1* cells, Dnm1 associated with the hyper-fused mitochondrial membrane, which is consistent with our observation that Fis1 localized to discrete foci in Δ*mdi1* cells (Fig. 3C, arrows). Thus, the defect in mitochondrial fission in the absence of Mdi1 is not due to loss of Dnm1 recruitment to mitochondria.

Given that Dnm1 can be recruited to mitochondria even in the absence of Mdi1, we asked how Dnm1 localization was affected during acutely induced mitochondrial division. We therefore treated cells expressing Dnm1-EGFP and mito-DsRed with CCCP and imaged by fluorescence microscopy. In wild type cells, fragments of mitochondria were often associated with Dnm1 foci (Fig. 3D, arrows). In contrast, in Δ*fis1* cells, Dnm1 failed to be recruited to mitochondria and often concentrated in bright focal structures in the cytosol (Fig. 3D). However, consistent with our observations in Δ*dnm1* cells (Fig. 3A), mitochondrial hyper-constriction upon CCCP treatment still occurred in the absence of Fis1. Remarkably, even though mitochondria are hyper-fused in the absence of either Fis1 or Mdi1, Dnm1 was able to assemble in focal structures on the hyper-constricted mitochondria only in Δ*mdi1* cells (Fig. 3D). Indeed, the mitochondria in these cells appeared so constricted they seem nearly fragmented, with Dnm1 commonly interspersed between each “bead” of mitochondrial signal at sites where the matrix marker signal appeared diminished (see Fig. 3D, linescan). To verify that mitochondrial division had not completed post-CCCP treatment in Δ*mdi1* cells when mitochondria appeared hyper-constricted, we co-expressed Dnm1-EGFP, mito-TagBFP (matrix), and Tom20-mCherry to label the OMM (Hughes et al., 2016), and imaged cells by confocal microscopy. Remarkably, even at sites where Dnm1 (green) was recruited to mitochondria and the matrix marker (mito-TagBFP; blue) appeared to be discontinuous, the OMM (Tom20-mCherry; magenta) appeared to remain connected (Fig. 3E, arrows). Together, these data indicate that Dnm1 recruitment to sites of constriction is insufficient to drive mitochondrial division to completion in the absence of Mdi1.

Based on the ability of Mdi1 to form foci independently of Dnm1 and our observation that Dnm1 is insufficient to complete CCCP-driven mitochondrial fission in the absence of Mdi1, we considered that the role of Mdi1 could be to divide the IMM independently. To test this possibility, we co-expressed chromosomally-tagged markers for both the OMM (Tom20-mCherry, magenta) and the IMM (Pam17-EGFP, green; (Wurm and Jakobs, 2006)) in wild type cells or in cells lacking Dnm1 or Mdi1. The expression of the tagged mitochondrial proteins did not negatively impact mitochondrial morphology, which appeared tubular as expected in wild type cells and net-like in the absence of either Dnm1 or Mdi1 (Fig. 3F). We then treated with CCCP for 45 minutes to induce mitochondrial hyper-constrictions in cells lacking either Dnm1 or Mdi1, which we could readily detect with Tom20-mCherry as thin threads between adjacent sections of mitochondria (Fig. 3G). We then asked whether the IMM marker, Pam17-EGFP, could be identified at these hyper constrictions, finding it was present in 100% of constrictions, both in the absence of Mdi1 and in Δ*dnm1* cells where Mdi1 was still present (Fig. 3G, see linescan). Thus, Mdi1 assembles within mitochondria independently of Dnm1, but likely does not independently mediate fission of the IMM.

### Mdi1 plays a functionally conserved role in mitochondrial division

Evolutionary analysis using CLIME (Li et al., 2014) and InterPro (Paysan-Lafosse et al., 2023) indicates that sequence homologs of Mdi1 are widely found within fungal species, though are notably absent from metazoans (Fig. 4A). To next determine if Mdi1 plays a similar role in mitochondrial division in other species, we sought to determine the effect of its loss in the fission yeast *Schizosaccharomyces pombe.* As in budding yeast, fission yeast mitochondria appear as part of a semi-continuous tubular network as assessed by microscopy of cells expressing the matrix marker mito-mCherry (Fig. 4B-4C). In wild type cells, mitochondria occasionally appeared more hyper-fused and contained smaller nets in ∼20% of cells. However, in the absence of mitochondrial fission (Δ*dnm1*), consistent with published observations (Dong et al., 2022), mitochondria formed extensive hyper-fused nets in the majority cells (Fig. 4B-4C). Occasionally the mitochondria in fission yeast Δ*dnm1* cells appeared aggregated or hyper-constricted, similar to budding yeast Δ*dnm1* cells that were treated with CCCP. In the absence of Mdi1, mitochondria regularly formed hyper-fused nets as in Δ*dnm1* cells, though also had abundant hyper-constricted, but interconnected, mitochondria (Fig. 4B-4C).

**Figure 4.**
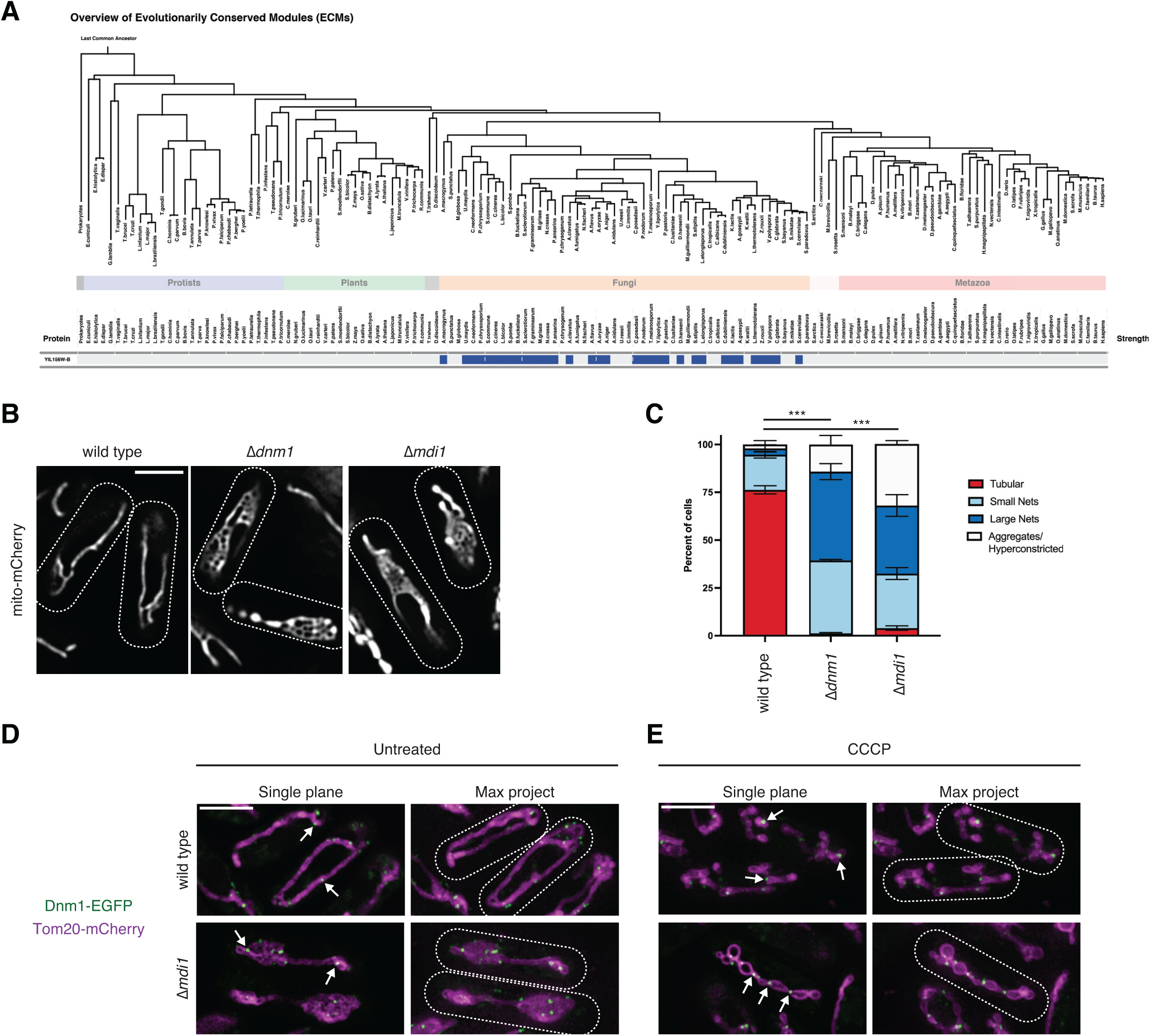
Mdi1 plays a functionally conserved role in mitochondrial division. **(A)** CLIME analysis (Li et al., 2014) of Mdi1 identifies conservation among fungal, but not plant or metazoan, species. **(B)** Single plane confocal fluorescence microscopy images of the indicated *S. pombe* strains expressing the matrix marker mito-mCherry. **(C)** A graph depicting categorization of mitochondrial morphology of cells as in (B). Data shown represent ∼100 cells per strain in each of three independent experiments and bars indicate S.E.M. Asterisks (***p<0.001) represent unpaired two-tailed *t* tests. **(D)** Single plane (left) and maximum intensity projection (right) confocal fluorescence microscopy images of wild type (top) and Δ*mdi1* (bottom) fission yeast cells expressing chromosomally tagged Tom20-mCherry (magenta) and Dnm1-EGFP (green). Arrows mark sites of Dnm1 recruitment to mitochondria. **(E)** As in (D) for cells treated for 30 minutes with 25 µM CCCP. Cell boundaries are indicated with dotted lines. Scale bars = 4 µm (B), 5 µm (D, E).

We next asked whether Dnm1 remains recruited to mitochondria in the absence of Mdi1 in fission yeast as it does in budding yeast. We generated endogenous chromosomally integrated fusions of Tom20-mCherry and Dnm1-EGFP in wild type and *Δmdi1* cells. As in mito-mCherry expressing cells, Tom20-mCherry labeled mitochondria appeared netlike in the absence of Mdi1 (Fig. 4D). Additionally, Dnm1-EGFP localized to discrete foci along the mitochondrial network in wild type cells, similar to budding yeast and consistent with published observations (Fig. 4D, arrows) (Jourdain et al., 2009). Comparable to our results in budding yeast, loss of Mdi1 did not impact Dnm1 recruitment to mitochondria and Dnm1 foci appeared associated with the hyper-fused net structures (Fig. 4D, arrows). Next, we induced fragmentation of the mitochondrial network with CCCP. Strikingly, in Δ*mdi1* cells, after CCCP treatment, Tom20-mCherry labeled mitochondria frequently had a hyper-constricted beads-on-a-string appearance and Dnm1 could be observed at discrete foci at each constriction site (Fig. 4E, arrows). Thus, our data indicates Mdi1 plays a conserved role as a pro-fission factor that is not required for Dnm1 recruitment but is required to facilitate the completion of mitochondrial division.

### A putative amphipathic alpha-helix is required for Mdi1 function

Analysis using InterPro indicates Mdi1 is defined by a domain of unknown function (DUF1748) (Paysan-Lafosse et al., 2023). AlphaFold 2 predictions of fungal DUF1748-containing proteins suggest they all contain two lengthy alpha-helices as in the case of Mdi1, although notably our data suggest that Mdi1 is not an integral membrane protein (Fig. S2C). We therefore examined structural characteristics of Mdi1 homologs to identify conserved motifs. Strikingly, we noticed that the second predicted α-helix appears amphipathic in helical wheel projection plots for each Mdi1 homolog we examined (Fig. 5A).

**Figure 5.**
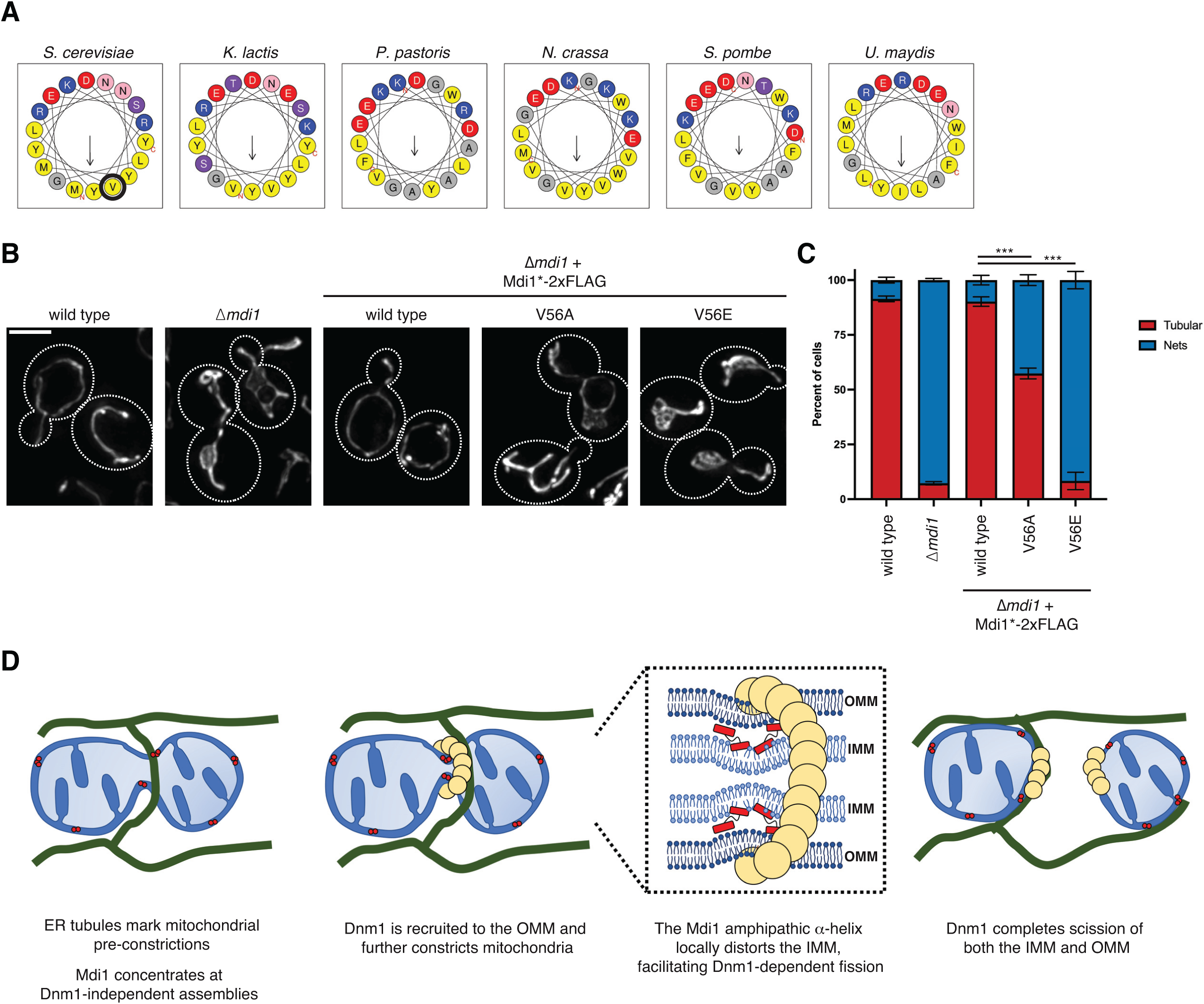
A putative amphipathic alpha-helix is required for Mdi1 function. **(A)** Helical wheel projection plots generated by HeliQuest (Gautier et al., 2008) of the second α-helical region of the indicated fungal orthologues of Mdi1. Valine 56 of *S. cerevisiae* Mdi1 is circled in black. (B) Maximum intensity projections of deconvolved epifluorescence microscopy images of the indicated yeast strains expressing mito-dsRed. Mdi1*(V56A)-2xFLAG and Mdi1*(V56E)-2xFLAG are driven by an estradiol-controlled GalL promoter and all strains were treated with 3 nM β-estradiol. (C) A graph depicting categorization of mitochondrial morphology of cells as in (B). Data shown represent at least 75 cells per strain in each of three independent experiments and bars indicate S.E.M. Asterisks (***p<0.001) represent unpaired two-tailed *t* tests. (D) A model for a potential role of Mdi1 in facilitating mitochondrial fission. Cell boundaries are indicated with dotted lines. Scale bar = 3 µm. See also Figure S3.

Amphipathic α-helices are common structural motifs that can promote membrane targeting of proteins. The hydrophobic side of the alpha helix can embed between hydrocarbon phospholipid tails, often causing lipid packing defects that have the capacity to bend or locally deform the membrane (Gimenez-Andres et al., 2018). To determine if the amphipathic character of the Mdi1 α-helix contributes to Mdi1 function, we generated mutations to budding yeast Mdi1, changing a valine residue on the hydrophobic side of the α-helix to a less bulky alanine (V56A) and to a charged glutamate (V56E) (Fig. 5A, circled in black). Expression of the Mdi1*-2xFLAG constructs with these mutations failed to express stably, and to counter this, we engineered a strain to drive their expression conditionally in the presence of estradiol so that they exceeded expression of wild type Mdi1*-2xFLAG controlled by its native promoter (Fig. S3A; (Veatch et al., 2009)).

Remarkably, these Mdi1 mutations led to defects in protein function as assayed by their ability to promote normal mitochondrial morphology. Unlike strains which express Mdi1*-2xFLAG, which appear to have normal mitochondrial morphology, cells expressing Mdi1*(V56A)-2xFLAG had net-like mitochondria in nearly half of cells and mitochondria in Mdi1*(V56E)-2xFLAG expressing cells appeared similar to those of Δ*mdi1* cells (Fig. 4B-4C). Because the Mdi1*(V56E) mutant failed to rescue mitochondrial morphology defects of Δ*mdi1* cells, we performed a protease protection assay and confirmed that the protein correctly targeted to the IMS (Fig. S3B). These data suggest that although the mutant protein localizes to the correct mitochondrial compartment, mitochondrial fission remains defective in these cells. In summary, our findings indicate that the amphipathic nature of a predicted Mdi1 α-helix is critical for Mdi1 to promote mitochondrial division.

## Discussion

While extensive work has been done to characterize Dnm1/Drp1 behavior in vitro and demonstrate its ability to utilize GTP hydrolysis to constrict membranes (Basu et al., 2017; Francy et al., 2015; Ingerman et al., 2005; Kalia et al., 2018; Kamerkar et al., 2018; Mears et al., 2011), it has remained unclear whether the proteins perform unassisted scission of both mitochondrial membranes. Our data indicate that the soluble, IMS-localized micropeptide Mdi1 plays a conserved role in facilitating Dnm1-dependent mitochondrial division in fungal species. Loss of Mdi1 in either budding yeast or fission yeast leads to mitochondrial hyper-fusion caused by an inability of Dnm1 to complete fission of the organelle, even despite its recruitment to hyper-constriction sites during induced division.

Mdi1 localizes to discrete focal structures within the IMS that can be spatially linked to sites of mitochondrial division, though it can concentrate at focal structures independently from Dnm1. However, Mdi1 is not required for Dnm1-independent mitochondrial pre-constriction, nor does it appear to promote IMM division independently from Dnm1. Mdi1 orthologs have a structurally conserved amphipathic α-helix, mutations in which prevent Mdi1 function in mitochondrial division, suggesting it may work in part by associating with and/or remodeling membranes. One model for a role for Mdi1 is that its amphipathic α-helix may allow it to bind and locally distort regions of the IMM and/or OMM, allowing Dnm1 to more easily constrict and simultaneously divide both the OMM and the IMM (Fig. 5D).

While Dnm1 is recruited to mitochondria via the OMM tail-anchored receptor Fis1, Fis1 has no functional domains exposed to the IMS, raising the question of how Mdi1 assemblies are positioned and associate with mitochondrial division sites. One possibility is that Mdi1 senses and assembles with specific phospholipids that enrich at discrete sites along the OMM or IMM, perhaps in proximity to contact sites with the endoplasmic reticulum. Alternatively, Mdi1 may have binding partners that are integral to either mitochondrial membrane that help position it near division sites. While our working model is that Mdi1 performs a direct role in division, it remains possible that Mdi1 may work indirectly via yet to be determined factors. Additionally, while our data suggests that Mdi1 cannot independently divide the IMM, we cannot rule out the possibility that it participates in a step of IMM scission that requires Dnm1-mediated hyper-constriction.

Our findings also beg the question of whether an IMS-localized factor works analogously to Mdi1 outside of fungal species. Some primitive eukaryotes such as the red algae *Cyanidioschyzon merolae* utilize the bacterial division machinery FtsZ to help facilitate mitochondrial division by constricting the IMM from within the matrix during Dnm1-mediated fission (Nishida et al., 2003; Osteryoung and Nunnari, 2003). Meanwhile, work from the Voeltz lab has suggested that in human cells, Drp1 cannot complete fission of mitochondria without subsequent activity of Dynamin 2 at division sites (Lee et al., 2016). Thus, it is possible that primitive eukaryotes, yeast, and humans each evolved to solve this issue through different mechanisms. However, Dynamin 2 is not universally required for mitochondrial division (Fonseca et al., 2019; Kamerkar et al., 2018), and Drp1 is capable of severing membrane tubules in vitro (Kamerkar et al., 2018). That said, it is not clear whether Drp1 is sufficient to completely divide two membrane bilayers in vivo. Additional micropeptides in humans are still being identified (Sandmann et al., 2023); for example, a matrix-targeted respiratory complex assembly factor is coincidentally encoded in the 5’-UTR of the mitochondrial fission protein MiD49/MIEF1 (Rathore et al., 2018). Thus, while Mdi1 is not conserved in human cells, structurally or functionally analogous proteins could conceivably work from inside mitochondria to cooperate in Drp1-mediated division in metazoans.

## Supplemental Data

**Figure S1.**
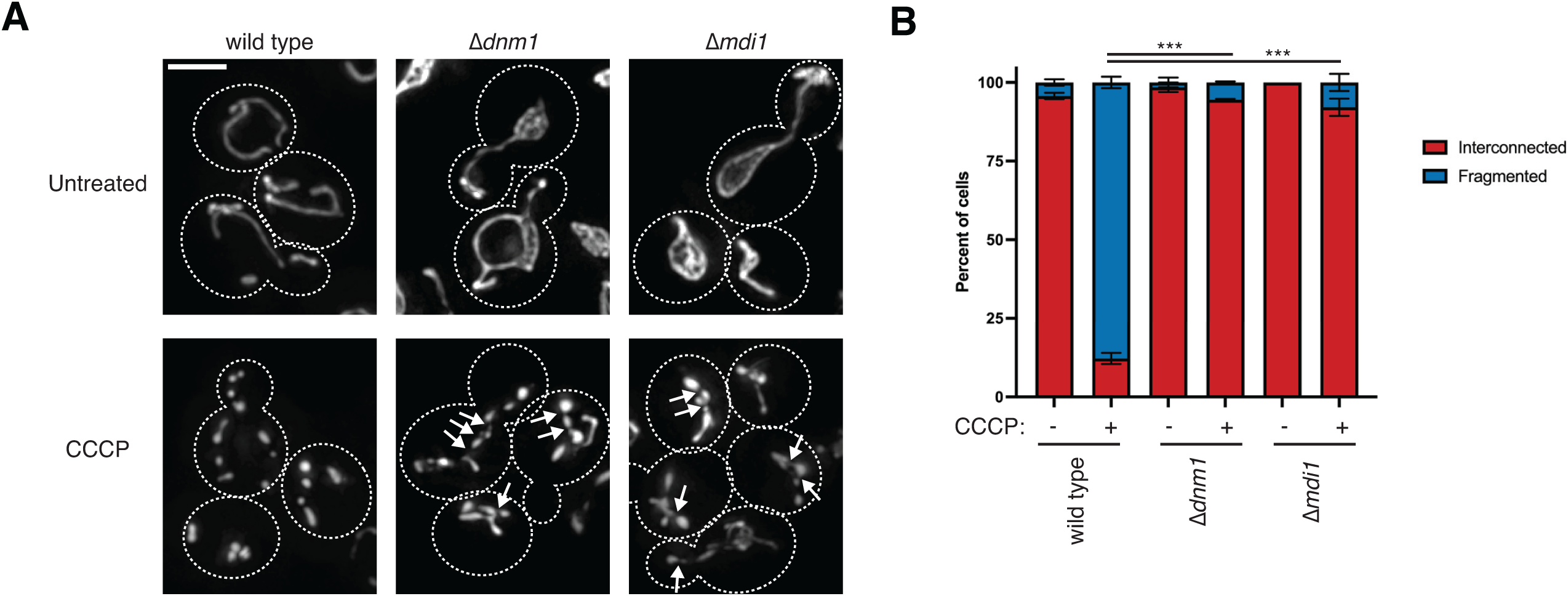
Mdi1 is required for CCCP-induced mitochondrial division. **(A)** Maximum intensity projections of deconvolved epifluorescence microscopy images of the indicated yeast strains expressing mito-dsRed before (top) or after (bottom) treatment with 25 µM CCCP for 45 minutes. Arrows mark mitochondrial hyper-constrictions that occur after CCCP treatment. **(B)** A graph depicting categorization of mitochondrial morphology of cells as in (A). Data shown represent at least 75 cells per strain in each of three independent experiments and bars indicate S.E.M. Asterisks (***p<0.001) represent unpaired two-tailed *t* tests. Cell boundaries are indicated with dotted lines. Scale bar = 3 µm.

**Figure S2.**
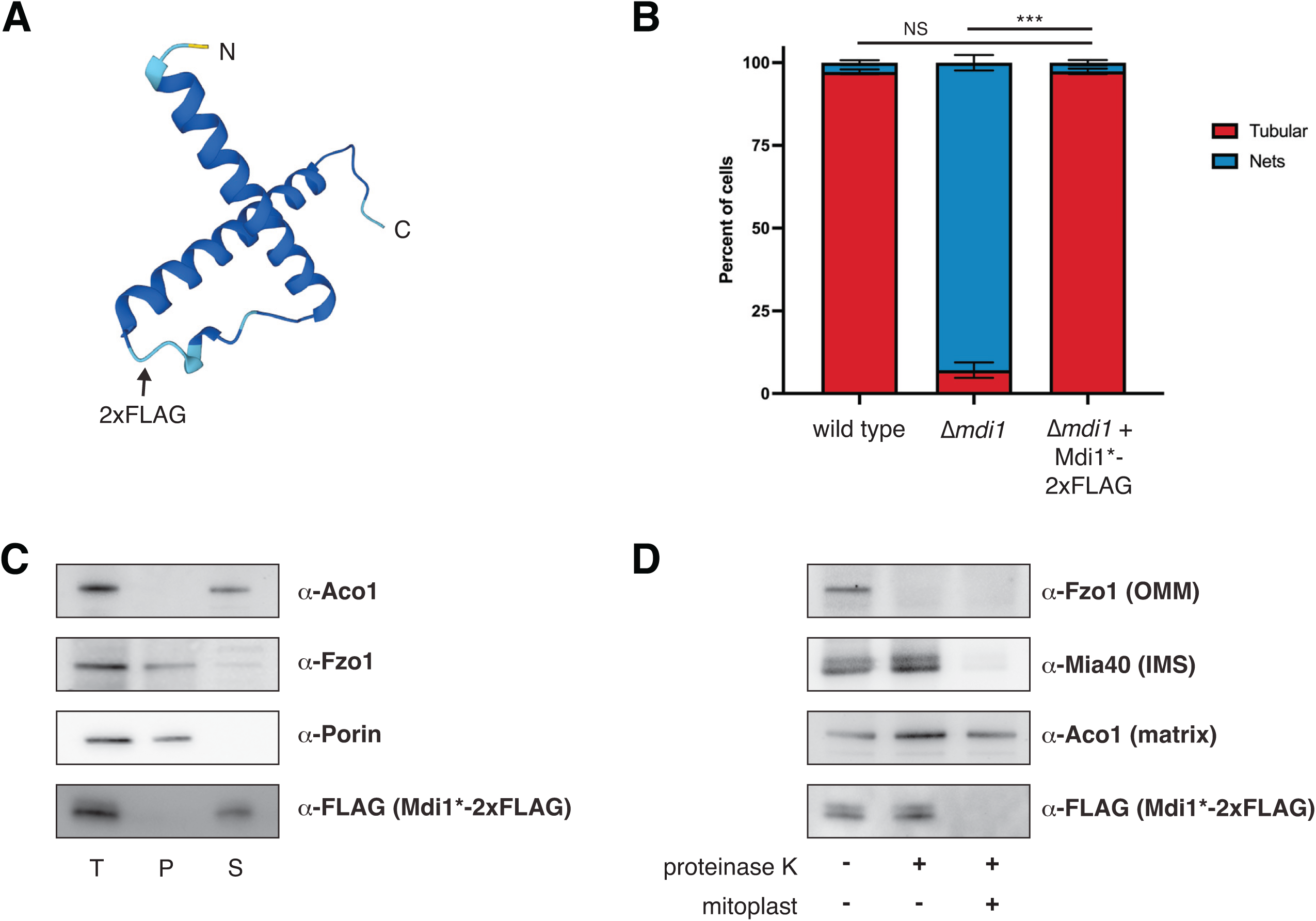
Mdi1 is a soluble intermembrane space protein. **(A)** AlphaFold 2 predicted structure of Mdi1 (Jumper et al., 2021; Varadi et al., 2022). Arrow marks site of insertion of internal 2xFLAG tag. **(B)** A graph depicting categorization of mitochondrial morphology of the indicated yeast strains expressing mito-dsRed. Data shown represent at least 75 cells per strain in each of three independent experiments and bars indicate S.E.M. Asterisks (***p<0.001) represent unpaired two-tailed *t* test. NS indicates not statistically significant. **(C)** Western analysis with the indicated antibodies of mitochondria isolated from an Δ*mdi1* strain expressing Mdi1*-2xFLAG and subjected to alkaline extraction. Crude membranes were incubated with 0.1M Na_2_CO_3_ pH11.5, and total (T), pellet (P), and supernatant (S) fractions were collected after centrifugation. **(D)** Western analysis with the indicated antibodies of mitochondria isolated from an Δ*mdi1* strain expressing Mdi1*-2xFLAG and subjected to protease protection analysis. Mitochondria were treated where indicated with proteinase K. Mitoplast sample indicates selective disruption of the OMM by a combination of osmotic swelling and mechanical disruption.

**Figure S3.**
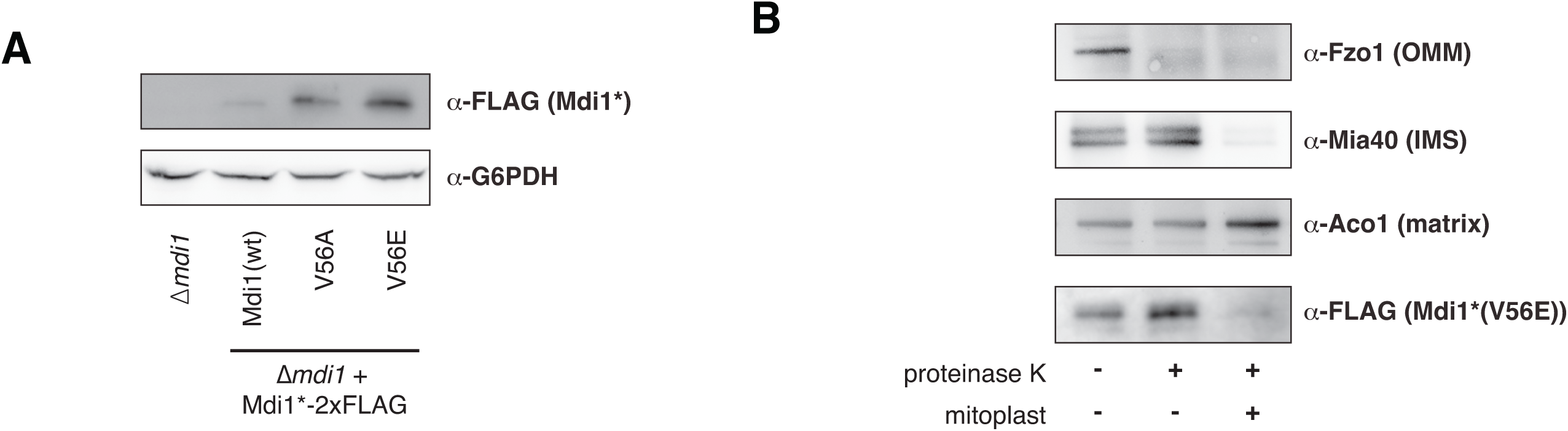
The Mdi1(V56E) mutation does not prevent targeting to the IMS. **(A)** Western analysis with the indicated antibodies of whole cell lysates of the indicated yeast strains. Expression of Mdi1*(V56A)-2xFLAG and Mdi1*(V56E)-2xFLAG were driven by an estradiol-controlled GalL promoter and all strains were treated with 3 nM β-estradiol. **(B)** Western analysis with the indicated antibodies of isolated mitochondria from an Δ*mdi1* strain expressing Mdi1*(V56E)-2xFLAG subjected to protease protection analysis. Mitochondria were treated where indicated with proteinase K. Mitoplast sample indicates selective disruption of the OMM by a combination of osmotic swelling and mechanical disruption.

## Materials and Methods

### Yeast growth

*Saccharomyces cerevisiae* strains were constructed in the W303 genetic background (*ade2-1*; *leu2-3*; *his3-11, 15*; *trp1-1*; *ura3-1*; *can1-100*). The *fzo1-1* and *fzo1-1* Δ*dnm1* yeast strains were a kind gift of Laura Lackner (Northwestern). Routine cell growth was performed in YPD (1% yeast extract, 2% peptone, 2% glucose), YPEG (1% yeast extract, 2% peptone, 3% ethanol, 3% glycerol), or synthetic complete dextrose (SCD; 2% glucose, 0.7% yeast nitrogen base, amino acids). Wild type haploid *Schizosaccharomyces pombe* was a kind gift of Mike Henne (UT Southwestern). All *S. pombe* strains were routinely grown in YES (0.5% yeast extract, 3% glucose, 225 mg/L adenine, 225mg/L leucine, 225 mg/L histidine, 225 mg/L uracil, 225 mg/L lysine) or as indicated in EMM (Sunrise Science; supplemented with 225 mg/L adenine, 225mg/L leucine, 225 mg/L histidine, 225 mg/L uracil, 225 mg/L lysine, 20 mg/L thiamine).

### Plasmids and yeast strain construction

All deletions in *S. cerevisiae* or *S. pombe* were made using PCR-based homologous recombination replacing the entire ORF with the HIS, NatMX6 or HphMX6 cassettes from pFA6a-series plasmids using lithium acetate transformation (Longtine et al., 1998). C-terminal protein fusions were integrated at the indicated endogenous chromosomal locus and driven by the native promoter, except where indicated for Mdi1, and were generated using pFA6a-link-yEGFP-*SpHIS5*, pFA6a-link-yEGFP-Kan, pFA6a-link-yEGFP-NatMX, pFA6a-mCherry-HphMX6 (see below), or pFA6a-mCherry-Kan (Lackner et al., 2013; Sheff and Thorn, 2004; Tirrell et al., 2020). pFA6a-mCherry-HphMX6 was generated by cloning the HphMX6 cassette into the BglII/EcoRV sites of pFA6a-mCherry-Kan, replacing the KanMX cassette.

The following plasmids were used to visualize the mitochondrial matrix in *S. cerevisiae*: pYX142 mito-dsRed (Friedman et al., 2011), pRS304 mito-DsRed, and pRS304 mito-TagBFP. pRS304 mito-DsRed was generated by cloning the mito-DsRed cassette from pRS305 mito-DsRed (pLL19; (Abrisch et al., 2020)) into the NotI/SacI sites of pRS304. pRS304 mito-TagBFP was generated by PCR amplifying mito-TagBFP from pVT100u-mt-TagBFP (Friedman et al., 2015) and cloning into the NcoI/AscI sites of pRS304 mito-DsRed, replacing mito-DsRed.

An expression plasmid of internally tagged Mdi1*-2xFLAG (pRS306 Mdi1(Leu37)-2xFLAG) was generated by isothermal assembly into the XhoI/NotI sites of pRS306 (Sikorski and Hieter, 1989) with DNA fragments consisting of the Mdi1 promoter, the Mdi1 coding sequence, an internal 2xFLAG tag after amino acid Leu37 flanked on each side by an additional glycine and serine, and the Mdi1 terminator.

To express mutants of Mdi1*-2xFLAG, the endogenous Mdi1 promoter of pRS306-Mdi1(Leu37)-2xFLAG was replaced with the GalL promoter that was PCR amplified from pYM-N29 (Janke et al., 2004), generating pRS306 GalLprom-Mdi1(Leu37)-2xFLAG. V56A and V56E mutations of Mdi1 were subsequently generated by site directed mutagenesis.

An expression plasmid of Mdi1-GFP11_x7_ was generated as follows. First, pFA6a-GFP11_x7_-Kan was generated by replacing the yEGFP cassette of pKT127 (Sheff and Thorn, 2004) with GFP11_x7_ amplified from pHRm-NLS-dCas9-GFP11x7-NLS (Addgene 70224; (Kamiyama et al., 2016). Then, Mdi1 was C-terminally tagged at the endogenous chromosomal locus with GFP11_x7_ preceded by a 24 amino acid linker by PCR-based homologous recombination as described above. Genomic DNA was then isolated from this strain and used to amplify the Mdi1 promoter, coding sequence, linker, GFP11_x7_, and terminator into the XhoI/NotI sites of pRS306 by isothermal assembly.

A GFP1-10 expression plasmid that co-expresses mito-DsRed, which enabled visualization of Mdi1-GFP11_x7_ (pRS304 Cyb2-GFP1-10 mito-DsRed), was generated by isothermal assembly of (1) the GPD promoter from pYM-N17 (Janke et al., 2004), (2) sequence coding for amino acids 1-167 of Cyb2, and (3) GFP1-10 from pFA6a-link-yGFP1-10-CaURA3MX (Addgene 86419; (Smoyer et al., 2016)) and subsequent cloning into the XhoI/NotI sites of pRS304 mito-DsRed.

An EGFP-Fis1 expression plasmid (pRS306 EGFP-Fis1) was generated by PCR amplifying (1) the Fis1 promoter, (2) the yEGFP coding sequence, and (3) the Fis1 coding sequence and terminator, and cloning into the XhoI/NotI sites of pRS306 by isothermal assembly.

Mdi1 and Fis1 expression plasmids were linearized and integrated into the yeast genome at the *ura3* locus. To control expression of mutated Mdi1 constructs driven by the GalL promoter with β-estradiol, pAGL-KanMX6 (Gok et al., 2022) was linearized and integrated at the *leu2* locus. Mitochondrial matrix targeted fluorescent proteins in budding yeast were either expressed from low copy plasmids maintained by auxotrophic selection or linearized and integrated into the *trp1* locus. Combinations of multiple tags and/or deletions were generated by strain crossing and tetrad dissection and/or by serial PCR-based homologous recombination. To visualize the mitochondrial matrix in *S. pombe*, mito-mCherry (mitoRED::Hyg; (Kraft and Lackner, 2019)) was linearized with NotI and introduced into the *leu1* locus.

### Fluorescence microscopy and analysis

All epifluorescence microscopy, where noted, was performed with a Nikon Eclipse Ti inverted epifluorescence microscope equipped with a Hamamatsu Orca-Fusion sCMOS camera and a Nikon 100x 1.45-NA objective and acquired with Nikon Elements. Z-series images were acquired with a 0.2 µm step size. All images were deconvolved using AutoQuant X3 (10 iterations, blind deconvolution, and low noise).

All confocal fluorescence microscopy, where noted, was performed on a Nikon Spinning Disk Confocal microscope with Yokogawa CSU-W1 SoRa and equipped with a Hamamatsu Orca-Fusion sCMOS camera and a Nikon 100x 1.45 NA objective. Z-series images were acquired with a 0.2 µm step size (except for timelapse microscopy as noted below) and the standard resolution disk with 50 µm pinholes.

Linear adjustments to images were made with ImageJ/Fiji. Linescan analysis was performed with slight modifications to the RGB Profiles Tool macro (https://imagej.nih.gov/ij/macros/tools/RGBProfilesTool.txt).

Strains were grown to exponential phase in SCD media with the appropriate auxotrophic selection (fission yeast were grown in EMM media), concentrated, and immobilized on a 3% agarose bed of growth media on cavity microscope slides. Multiple fields of view were imaged per strain or condition in each of at least 3 independent experiments. Except where indicated below, cells were grown at 30°C and imaged at room temperature. For experiments involving *fzo1-1* strains, cells were grown to exponential phase at room temperature and imaged at room temperature, or shifted to 37°C for 20 minutes and imaged at 37°C. For CCCP treatment, cells were grown to exponential phase at 30°C, media was supplemented with CCCP (25 µM; Sigma C2759) and cells were allowed to continue growth at 30°C for the indicated times prior to imaging at room temperature. For expression of Mdi1 point mutants, standard imaging analysis was performed for cells grown constitutively in the presence of β-estradiol (3nM; Calbiochem 3301).

To assess mitochondrial morphology, images were blinded prior to analysis, cells were manually categorized as indicated in figures, and the number of cells analyzed per experiment is indicated in the associated figure legend. Statistical comparison was performed between each of the indicated samples by unpaired two-tailed t-test with Welch’s correction of the tubular or interconnected mitochondrial morphology category.

To characterize Mdi1 localization to mitochondrial division sites, 3 images centered on the midplane of cells were captured in 0.4µm steps at 10 second intervals for 2 minutes. Maximum intensity projections were generated in ImageJ and mitochondrial division sites were identified with mito-dsRed blind to Mdi1-GFP11_x7_/IMS-GFP1-10 signal. Mdi1 focal localization relative to the division site was then manually assessed.

To assess IMM connectivity at hyper-constricted mitochondria, maximum intensity projections of images of CCCP-treated cells co-expressing Pam17-EGFP (IMM) and Tom20-mCherry (OMM) were generated using ImageJ. Tom20-marked mitochondrial hyper-constrictions were identified blind to Pam17-EGFP signal, and then manually assessed for continuity of Pam17 fluorescence at the constriction site.

### Cell growth analysis

For analysis of growth of yeast cells on ethanol/glycerol media, assays were performed by growing cells to exponential phase in YPD at room temperature, pelleting, and resuspending cells in water at a concentration of 0.5 OD600/mL. 5µL of 10-fold serial dilutions of cells were plated on YPEG plates and incubated at the indicated temperatures.

### Mitochondria isolation, alkaline extraction, and protease protection analysis

Mitochondria were isolated by differential centrifugation as previously described (Meeusen et al., 2004), with the following modifications. Briefly, 500-1000 ODs of yeast cells were grown to exponential phase in YPEG media (Mdi1*(V56E)-2xFLAG was grown in the presence of 1nM estradiol). After harvesting cells and washing with water, spheroplasts were generated by digesting cell walls with 4mg/ml zymolyase 20T (Sunrise Science) in 1.2M sorbitol. Spheroplasts were washed and resuspended in a minimal volume of cold MIB (0.6M sorbitol, 50mM KCl, 20mM HEPES pH7.4) and homogenized with a tight fitting dounce. Un-lysed cells and large debris were removed by low speed centrifugation (2400g, 5 minutes, 4°C). Crude mitochondria were enriched by centrifugation of the supernatant (13000g, 15 minutes, 4°C). The mitochondrial pellet was resuspended in a minimal volume of cold MIB to a final concentration of 5-10 mg/ml. Protein concentration was measured by a Bradford assay (BioRad) and 200 µg aliquots were flash frozen in liquid nitrogen and stored at -80°C.

Alkaline extraction to determine solubility of Mdi1 was performed as previously described, with minor modifications (Hoppins et al., 2011). Equivalent amounts of crude mitochondria were washed, and pellets were either directly resuspended in Laemmli sample buffer or resuspended in 0.1 M Na_2_CO_3_ pH11.5 and incubated 30 minutes on ice. Na_2_CO_3_-treated mitochondria were subjected to centrifugation (60 minutes, 100000g, 4°C) and the pellet was resuspended in an equivalent amount of Laemmli sample buffer. Supernatant proteins were precipitated with TCA (12.5%, 30 minutes on ice), pelleted by centrifugation (10 minutes, 16,000g, 4°C), washed with cold acetone, and resuspended in Laemmli sample buffer. Samples were analyzed by SDS-PAGE and Western blot as described below.

Protease protection analysis of Mdi1 was performed as described previously with minor modifications (Hoppins et al., 2011). Crude mitochondria were washed with MIB, distributed into 3 equivalent tubes, pelleted, and resuspended in equivalent volumes as follows: two tubes were resuspended in MIB buffer and one tube was resuspended in mitoplast buffer (20mM HEPES pH7.4). Mitoplast samples were incubated on ice for 15 minutes and then mechanically disrupted by pipetting up and down 15 times. MIB-resuspended mitochondria were mock treated or treated with proteinase K (100µg/ml), mitoplast samples were treated with proteinase K, and all samples were incubated on ice an additional 15 minutes. 2mM PMSF was added to all samples to stop the protease reaction and incubated 5 minutes on ice. Samples were subjected to centrifugation (10,400g, 15 minutes, 4°C), pellets were resuspended in MIB with 1x protease inhibitor cocktail (MilliporeSigma 539131), and precipitated with TCA as above. Samples were resuspended in Laemmli sample buffer and analyzed by SDS-PAGE/Western blotting as described below.

### Whole cell extracts and western analysis

For whole cell extracts, cells were grown to exponential phase in SCD. 0.25 OD600 cells were pelleted, washed with dH20, and extracts were prepared by alkaline extraction (0.255M NaOH, 1% 2-mercaptoethanol) followed by precipitation in 9% trichloroacetic acid. Precipitates were washed with acetone, dried, and resuspended in 50µL Laemmli protein sample buffer prior to Western analysis.

Whole cell lysates or crude mitochondria extracts were freshly prepared and incubated at 70°C (to enable optimal detection of Mdi1*-2xFLAG) or 95°C for 10 minutes prior to SDS-PAGE, transferred to 0.2µm pore size PVDF membranes, and immunoblotted with the following primary antibodies: mouse α-FLAG (1:1000, Sigma F1804), rabbit α-G6PDH (1:2000, Sigma A9521), mouse α-Porin (1:2000; ThermoFisher 459500), rabbit α-Fzo1 (1:1000, a kind gift of Jodi Nunnari, Altos Labs), α-Aco1 (1:10000, a kind gift of Anju Sreelatha, UT Southwestern), α-Mia40 (1:10000, a kind gift of Anju Sreelatha, UT Southwestern). To detect Mdi1*-2xFLAG, goat anti-rabbit HRP (Sigma) was used (1:10000) and signal was visualized with SuperSignal West Femto Substrate (ThermoFisher). All other proteins were detected with secondary antibodies conjugated to DyLight800 (1:10000, ThermoFisher). Images were acquired with a ChemiDoc MP Imaging System (BioRad). Linear adjustments to images were made with Photoshop (Adobe).

## Acknowledgements

We thank Laura Lackner (Northwestern) for critical reading of the manuscript, helpful discussions, and kindly providing yeast strains. We thank Mike Henne (UT Southwestern) and Maya Schuldiner (Weizmann Institute of Science) for helpful discussions. We thank Jodi Nunnari (Altos Labs) and Anju Sreelatha (UT Southwestern) for kindly providing antibodies. The UT Southwestern Quantitative Light Microscopy Facility, which is supported in part by NIH P30CA142543, provided access to the Nikon SoRa microscope (purchased with NIH 1S10OD028630-01 to Kate Luby-Phelps) and deconvolution software. This work was supported by a grant from the NIH to JF (R35GM137894). The authors declare no competing financial interests.

## Notes

### Competing Interest Statement

The authors have declared no competing interest.

## References

Abrisch, R.G., S.C. Gumbin, B.T. Wisniewski, L.L. Lackner, and G.K. Voeltz. 2020. Fission and fusion machineries converge at ER contact sites to regulate mitochondrial morphology. J Cell Biol. 219.

Anand, R., T. Wai, M.J. Baker, N. Kladt, A.C. Schauss, E. Rugarli, and T. Langer. 2014. The i-AAA protease YME1L and OMA1 cleave OPA1 to balance mitochondrial fusion and fission. J Cell Biol. 204:919–929.

Basu, K., D. Lajoie, T. Aumentado-Armstrong, J. Chen, R.I. Koning, B. Bossy, M. Bostina, A. Sik, E. Bossy-Wetzel, and I. Rouiller. 2017. Molecular mechanism of DRP1 assembly studied in vitro by cryo-electron microscopy. PLoS One. 12:e0179397.

Beasley, E.M., S. Muller, and G. Schatz. 1993. The signal that sorts yeast cytochrome b2 to the mitochondrial intermembrane space contains three distinct functional regions. EMBO J. 12:2303–2311.

Bleazard, W., J.M. McCaffery, E.J. King, S. Bale, A. Mozdy, Q. Tieu, J. Nunnari, and J.M. Shaw. 1999. The dynamin-related GTPase Dnm1 regulates mitochondrial fission in yeast. Nat Cell Biol. 1:298–304.

Bosch, J.A., B. Ugur, I. Pichardo-Casas, J. Rabasco, F. Escobedo, Z. Zuo, B. Brown, S. Celniker, D.A. Sinclair, H.J. Bellen, and N. Perrimon. 2022. Two neuronal peptides encoded from a single transcript regulate mitochondrial complex III in Drosophila. Elife. 11.

Chacinska, A., S. Pfannschmidt, N. Wiedemann, V. Kozjak, L.K. Sanjuan Szklarz, A. Schulze-Specking, K.N. Truscott, B. Guiard, C. Meisinger, and N. Pfanner. 2004. Essential role of Mia40 in import and assembly of mitochondrial intermembrane space proteins. EMBO J. 23:3735–3746.

Chakrabarti, R., W.K. Ji, R.V. Stan, J. de Juan Sanz, T.A. Ryan, and H.N. Higgs. 2018. INF2-mediated actin polymerization at the ER stimulates mitochondrial calcium uptake, inner membrane constriction, and division. J Cell Biol. 217:251–268.

Cho, B., H.M. Cho, Y. Jo, H.D. Kim, M. Song, C. Moon, H. Kim, K. Kim, H. Sesaki, I.J. Rhyu, H. Kim, and W. Sun. 2017. Constriction of the mitochondrial inner compartment is a priming event for mitochondrial division. Nat Commun. 8:15754.

Chu, Q., T.F. Martinez, S.W. Novak, C.J. Donaldson, D. Tan, J.M. Vaughan, T. Chang, J.K. Diedrich, L. Andrade, A. Kim, T. Zhang, U. Manor, and A. Saghatelian. 2019. Regulation of the ER stress response by a mitochondrial microprotein. Nat Commun. 10:4883.

Dong, F., M. Zhu, F. Zheng, and C. Fu. 2022. Mitochondrial fusion and fission are required for proper mitochondrial function and cell proliferation in fission yeast. FEBS J. 289:262–278.

Fonseca, T.B., A. Sanchez-Guerrero, I. Milosevic, and N. Raimundo. 2019. Mitochondrial fission requires DRP1 but not dynamins. Nature. 570:E34–E42.

Francy, C.A., F.J. Alvarez, L. Zhou, R. Ramachandran, and J.A. Mears. 2015. The mechanoenzymatic core of dynamin-related protein 1 comprises the minimal machinery required for membrane constriction. J Biol Chem. 290:11692–11703.

Friedman, J.R., L.L. Lackner, M. West, J.R. DiBenedetto, J. Nunnari, and G.K. Voeltz. 2011. ER tubules mark sites of mitochondrial division. Science. 334:358–362.

Friedman, J.R., A. Mourier, J. Yamada, J.M. McCaffery, and J. Nunnari. 2015. MICOS coordinates with respiratory complexes and lipids to establish mitochondrial inner membrane architecture. Elife. 4.

Friedman, J.R., and J. Nunnari. 2014. Mitochondrial form and function. Nature. 505:335–343.

Gao, S., and J. Hu. 2021. Mitochondrial Fusion: The Machineries In and Out. Trends Cell Biol. 31:62–74.

Gautier, R., D. Douguet, B. Antonny, and G. Drin. 2008. HELIQUEST: a web server to screen sequences with specific alpha-helical properties. Bioinformatics. 24:2101–2102.

Gimenez-Andres, M., A. Copic, and B. Antonny. 2018. The Many Faces of Amphipathic Helices. Biomolecules. 8.

Gok, M.O., N.O. Speer, W.M. Henne, and J.R. Friedman. 2022. ER-localized phosphatidylethanolamine synthase plays a conserved role in lipid droplet formation. Mol Biol Cell. 33:ar11.

Guo, Y., D. Li, S. Zhang, Y. Yang, J.J. Liu, X. Wang, C. Liu, D.E. Milkie, R.P. Moore, U.S. Tulu, D.P. Kiehart, J. Hu, J. Lippincott-Schwartz, E. Betzig, and D. Li. 2018. Visualizing Intracellular Organelle and Cytoskeletal Interactions at Nanoscale Resolution on Millisecond Timescales. Cell. 175:1430–1442 e1417.

Hermann, G.J., J.W. Thatcher, J.P. Mills, K.G. Hales, M.T. Fuller, J. Nunnari, and J.M. Shaw. 1998. Mitochondrial fusion in yeast requires the transmembrane GTPase Fzo1p. The Journal of Cell Biology. 143:359–373.

Hoppins, S., S.R. Collins, A. Cassidy-Stone, E. Hummel, R.M. Devay, L.L. Lackner, B. Westermann, M. Schuldiner, J.S. Weissman, and J. Nunnari. 2011. A mitochondrial-focused genetic interaction map reveals a scaffold-like complex required for inner membrane organization in mitochondria. The Journal of Cell Biology. 195:323–340.

Hughes, A.L., C.E. Hughes, K.A. Henderson, N. Yazvenko, and D.E. Gottschling. 2016. Selective sorting and destruction of mitochondrial membrane proteins in aged yeast. Elife. 5.

Hung, V., P. Zou, H.W. Rhee, N.D. Udeshi, V. Cracan, T. Svinkina, S.A. Carr, V.K. Mootha, and A.Y. Ting. 2014. Proteomic mapping of the human mitochondrial intermembrane space in live cells via ratiometric APEX tagging. Mol Cell. 55:332–341.

Ingerman, E., E.M. Perkins, M. Marino, J.A. Mears, J.M. McCaffery, J.E. Hinshaw, and J. Nunnari. 2005. Dnm1 forms spirals that are structurally tailored to fit mitochondria. The Journal of Cell Biology. 170:1021–1027.

Ishihara, N., M. Nomura, A. Jofuku, H. Kato, S.O. Suzuki, K. Masuda, H. Otera, Y. Nakanishi, I. Nonaka, Y. Goto, N. Taguchi, H. Morinaga, M. Maeda, R. Takayanagi, S. Yokota, and K. Mihara. 2009. Mitochondrial fission factor Drp1 is essential for embryonic development and synapse formation in mice. Nat Cell Biol. 11:958–966.

Janke, C., M.M. Magiera, N. Rathfelder, C. Taxis, S. Reber, H. Maekawa, A. Moreno-Borchart, G. Doenges, E. Schwob, E. Schiebel, and M. Knop. 2004. A versatile toolbox for PCR-based tagging of yeast genes: new fluorescent proteins, more markers and promoter substitution cassettes. Yeast. 21:947–962.

Jourdain, I., Y. Gachet, and J.S. Hyams. 2009. The dynamin related protein Dnm1 fragments mitochondria in a microtubule-dependent manner during the fission yeast cell cycle. Cell Motil Cytoskeleton. 66:509–523.

Jumper, J., R. Evans, A. Pritzel, T. Green, M. Figurnov, O. Ronneberger, K. Tunyasuvunakool, R. Bates, A. Zidek, A. Potapenko, A. Bridgland, C. Meyer, S.A.A. Kohl, A.J. Ballard, A. Cowie, B. Romera-Paredes, S. Nikolov, R. Jain, J. Adler, T. Back, S. Petersen, D. Reiman, E. Clancy, M. Zielinski, M. Steinegger, M. Pacholska, T. Berghammer, S. Bodenstein, D. Silver, O. Vinyals, A.W. Senior, K. Kavukcuoglu, P. Kohli, and D. Hassabis. 2021. Highly accurate protein structure prediction with AlphaFold. Nature. 596:583–589.

Kalia, R., R.Y. Wang, A. Yusuf, P.V. Thomas, D.A. Agard, J.M. Shaw, and A. Frost. 2018. Structural basis of mitochondrial receptor binding and constriction by DRP1. Nature. 558:401–405.

Kall, L., A. Krogh, and E.L. Sonnhammer. 2004. A combined transmembrane topology and signal peptide prediction method. J Mol Biol. 338:1027–1036.

Kamerkar, S.C., F. Kraus, A.J. Sharpe, T.J. Pucadyil, and M.T. Ryan. 2018. Dynamin-related protein 1 has membrane constricting and severing abilities sufficient for mitochondrial and peroxisomal fission. Nat Commun. 9:5239.

Kamiyama, D., S. Sekine, B. Barsi-Rhyne, J. Hu, B. Chen, L.A. Gilbert, H. Ishikawa, M.D. Leonetti, W.F. Marshall, J.S. Weissman, and B. Huang. 2016. Versatile protein tagging in cells with split fluorescent protein. Nat Commun. 7:11046.

Klecker, T., M. Wemmer, M. Haag, A. Weig, S. Bockler, T. Langer, J. Nunnari, and B. Westermann. 2015. Interaction of MDM33 with mitochondrial inner membrane homeostasis pathways in yeast. Sci Rep. 5:18344.

Kleele, T., T. Rey, J. Winter, S. Zaganelli, D. Mahecic, H. Perreten Lambert, F.P. Ruberto, M. Nemir, T. Wai, T. Pedrazzini, and S. Manley. 2021. Distinct fission signatures predict mitochondrial degradation or biogenesis. Nature. 593:435–439.

Korobova, F., V. Ramabhadran, and H.N. Higgs. 2013. An actin-dependent step in mitochondrial fission mediated by the ER-associated formin INF2. Science. 339:464–467.

Kraft, L.M., and L.L. Lackner. 2019. A conserved mechanism for mitochondria-dependent dynein anchoring. Mol Biol Cell. 30:691–702.

Kraus, F., K. Roy, T.J. Pucadyil, and M.T. Ryan. 2021. Function and regulation of the divisome for mitochondrial fission. Nature. 590:57–66.

Krogh, A., B. Larsson, G. von Heijne, and E.L. Sonnhammer. 2001. Predicting transmembrane protein topology with a hidden Markov model: application to complete genomes. J Mol Biol. 305:567–580.

Labrousse, A.M., M.D. Zappaterra, D.A. Rube, and A.M. van der Bliek. 1999. C. elegans dynamin-related protein DRP-1 controls severing of the mitochondrial outer membrane. Mol Cell. 4:815–826.

Lackner, L.L., H. Ping, M. Graef, A. Murley, and J. Nunnari. 2013. Endoplasmic reticulum-associated mitochondria-cortex tether functions in the distribution and inheritance of mitochondria. Proc Natl Acad Sci USA. 110:E458–467.

Lee, J.E., L.M. Westrate, H. Wu, C. Page, and G.K. Voeltz. 2016. Multiple dynamin family members collaborate to drive mitochondrial division. Nature. 540:139–143.

Legesse-Miller, A., R.H. Massol, and T. Kirchhausen. 2003. Constriction and Dnm1p recruitment are distinct processes in mitochondrial fission. Mol Biol Cell. 14:1953–1963.

Lewis, S.C., L.F. Uchiyama, and J. Nunnari. 2016. ER-mitochondria contacts couple mtDNA synthesis with mitochondrial division in human cells. Science. 353:aaf5549.

Li, Y., S.E. Calvo, R. Gutman, J.S. Liu, and V.K. Mootha. 2014. Expansion of biological pathways based on evolutionary inference. Cell. 158:213–225.

Longtine, M.S., A. McKenzie, D.J. Demarini, N.G. Shah, A. Wach, A. Brachat, P. Philippsen, and J.R. Pringle. 1998. Additional modules for versatile and economical PCR-based gene deletion and modification in Saccharomyces cerevisiae. Yeast. 14:953–961.

Manor, U., S. Bartholomew, G. Golani, E. Christenson, M. Kozlov, H. Higgs, J. Spudich, and J. Lippincott-Schwartz. 2015. A mitochondria-anchored isoform of the actin-nucleating spire protein regulates mitochondrial division. Elife. 4.

Mears, J.A., L.L. Lackner, S. Fang, E. Ingerman, J. Nunnari, and J.E. Hinshaw. 2011. Conformational changes in Dnm1 support a contractile mechanism for mitochondrial fission. Nat Struct Mol Biol. 18:20–26.

Meeusen, S., J.M. McCaffery, and J. Nunnari. 2004. Mitochondrial fusion intermediates revealed in vitro. Science. 305:1747–1752.

Messerschmitt, M., S. Jakobs, F. Vogel, S. Fritz, K.S. Dimmer, W. Neupert, and B. Westermann. 2003. The inner membrane protein Mdm33 controls mitochondrial morphology in yeast. The Journal of Cell Biology. 160:553–564.

Morgenstern, M., S.B. Stiller, P. Lubbert, C.D. Peikert, S. Dannenmaier, F. Drepper, U. Weill, P. Hoss, R. Feuerstein, M. Gebert, M. Bohnert, M. van der Laan, M. Schuldiner, C. Schutze, S. Oeljeklaus, N. Pfanner, N. Wiedemann, and B. Warscheid. 2017. Definition of a High-Confidence Mitochondrial Proteome at Quantitative Scale. Cell reports. 19:2836–2852.

Mozdy, A.D., J.M. McCaffery, and J.M. Shaw. 2000. Dnm1p GTPase-mediated mitochondrial fission is a multi-step process requiring the novel integral membrane component Fis1p. The Journal of Cell Biology. 151:367–380.

Nguyen, T.T., and G.K. Voeltz. 2022. An ER phospholipid hydrolase drives ER-associated mitochondrial constriction for fission and fusion. Elife. 11.

Nishida, K., M. Takahara, S.Y. Miyagishima, H. Kuroiwa, M. Matsuzaki, and T. Kuroiwa. 2003. Dynamic recruitment of dynamin for final mitochondrial severance in a primitive red alga. Proc Natl Acad Sci U S A. 100:2146–2151.

Nunnari, J., W.F. Marshall, A. Straight, A. Murray, J.W. Sedat, and P. Walter. 1997. Mitochondrial transmission during mating in Saccharomyces cerevisiae is determined by mitochondrial fusion and fission and the intramitochondrial segregation of mitochondrial DNA. Mol Biol Cell. 8:1233–1242.

Osman, C., T.R. Noriega, V. Okreglak, J.C. Fung, and P. Walter. 2015. Integrity of the yeast mitochondrial genome, but not its distribution and inheritance, relies on mitochondrial fission and fusion. Proc Natl Acad Sci U S A. 112:E947–956.

Osteryoung, K.W., and J. Nunnari. 2003. The division of endosymbiotic organelles. Science. 302:1698–1704.

Paysan-Lafosse, T., M. Blum, S. Chuguransky, T. Grego, B.L. Pinto, G.A. Salazar, M.L. Bileschi, P. Bork, A. Bridge, L. Colwell, J. Gough, D.H. Haft, I. Letunic, A. Marchler-Bauer, H. Mi, D.A. Natale, C.A. Orengo, A.P. Pandurangan, C. Rivoire, C.J.A. Sigrist, I. Sillitoe, N. Thanki, P.D. Thomas, S.C.E. Tosatto, C.H. Wu, and A. Bateman. 2023. InterPro in 2022. Nucleic Acids Res. 51:D418–D427.

Pickles, S., P. Vigie, and R.J. Youle. 2018. Mitophagy and Quality Control Mechanisms in Mitochondrial Maintenance. Curr Biol. 28:R170–R185.

Quintana-Cabrera, R., and L. Scorrano. 2023. Determinants and outcomes of mitochondrial dynamics. Mol Cell. 83:857–876.

Rath, S., R. Sharma, R. Gupta, T. Ast, C. Chan, T.J. Durham, R.P. Goodman, Z. Grabarek, M.E. Haas, W.H.W. Hung, P.R. Joshi, A.A. Jourdain, S.H. Kim, A.V. Kotrys, S.S. Lam, J.G. McCoy, J.D. Meisel, M. Miranda, A. Panda, A. Patgiri, R. Rogers, S. Sadre, H. Shah, O.S. Skinner, T.L. To, M.A. Walker, H. Wang, P.S. Ward, J. Wengrod, C.C. Yuan, S.E. Calvo, and V.K. Mootha. 2021. MitoCarta3.0: an updated mitochondrial proteome now with sub-organelle localization and pathway annotations. Nucleic Acids Res. 49:D1541–D1547.

Rathore, A., Q. Chu, D. Tan, T.F. Martinez, C.J. Donaldson, J.K. Diedrich, J.R. Yates, 3rd, and A. Saghatelian. 2018. MIEF1 Microprotein Regulates Mitochondrial Translation. Biochemistry. 57:5564-5575.

Rensvold, J.W., E. Shishkova, Y. Sverchkov, I.J. Miller, A. Cetinkaya, A. Pyle, M. Manicki, D.R. Brademan, Y. Alanay, J. Raiman, A. Jochem, P.D. Hutchins, S.R. Peters, V. Linke, K.A. Overmyer, A.Z. Salome, A.S. Hebert, C.E. Vincent, N.W. Kwiecien, M.J.P. Rush, M.S. Westphall, M. Craven, N.A. Akarsu, R.W. Taylor, J.J. Coon, and D.J. Pagliarini. 2022. Defining mitochondrial protein functions through deep multiomic profiling. Nature. 606:382–388.

Rhee, H.W., P. Zou, N.D. Udeshi, J.D. Martell, V.K. Mootha, S.A. Carr, and A.Y. Ting. 2013. Proteomic mapping of mitochondria in living cells via spatially restricted enzymatic tagging. Science. 339:1328–1331.

Sandmann, C.L., J.F. Schulz, J. Ruiz-Orera, M. Kirchner, M. Ziehm, E. Adami, M. Marczenke, A. Christ, N. Liebe, J. Greiner, A. Schoenenberger, M.B. Muecke, N. Liang, R.L. Moritz, Z. Sun, E.W. Deutsch, M. Gotthardt, J.M. Mudge, J.R. Prensner, T.E. Willnow, P. Mertins, S. van Heesch, and N. Hubner. 2023. Evolutionary origins and interactomes of human, young microproteins and small peptides translated from short open reading frames. Mol Cell. 83:994–1011 e1018.

Sesaki, H., and R.E. Jensen. 1999. Division versus fusion: Dnm1p and Fzo1p antagonistically regulate mitochondrial shape. The Journal of Cell Biology. 147:699–706.

Sheff, M.A., and K.S. Thorn. 2004. Optimized cassettes for fluorescent protein tagging in Saccharomyces cerevisiae. Yeast. 21:661–670.

Sikorski, R.S., and P. Hieter. 1989. A system of shuttle vectors and yeast host strains designed for efficient manipulation of DNA in Saccharomyces cerevisiae. Genetics. 122:19–27.

Smoyer, C.J., S.S. Katta, J.M. Gardner, L. Stoltz, S. McCroskey, W.D. Bradford, M. McClain, S.E. Smith, B.D. Slaughter, J.R. Unruh, and S.L. Jaspersen. 2016. Analysis of membrane proteins localizing to the inner nuclear envelope in living cells. J Cell Biol. 215:575–590.

Stefely, J.A., N.W. Kwiecien, E.C. Freiberger, A.L. Richards, A. Jochem, M.J.P. Rush, A. Ulbrich, K.P. Robinson, P.D. Hutchins, M.T. Veling, X. Guo, Z.A. Kemmerer, K.J. Connors, E.A. Trujillo, J. Sokol, H. Marx, M.S. Westphall, A.S. Hebert, D.J. Pagliarini, and J.J. Coon. 2016. Mitochondrial protein functions elucidated by multi-omic mass spectrometry profiling. Nat Biotechnol. 34:1191–1197.

Tharakan, R., and A. Sawa. 2021. Minireview: Novel Micropeptide Discovery by Proteomics and Deep Sequencing Methods. Front Genet. 12:651485.

Tieu, Q., and J. Nunnari. 2000. Mdv1p is a WD repeat protein that interacts with the dynamin-related GTPase, Dnm1p, to trigger mitochondrial division. The Journal of Cell Biology. 151:353-366.

Tirrell, P.S., K.N. Nguyen, K. Luby-Phelps, and J.R. Friedman. 2020. MICOS subcomplexes assemble independently on the mitochondrial inner membrane in proximity to ER contact sites. J Cell Biol. 219.

Varadi, M., S. Anyango, M. Deshpande, S. Nair, C. Natassia, G. Yordanova, D. Yuan, O. Stroe, G. Wood, A. Laydon, A. Zidek, T. Green, K. Tunyasuvunakool, S. Petersen, J. Jumper, E. Clancy, R. Green, A. Vora, M. Lutfi, M. Figurnov, A. Cowie, N. Hobbs, P. Kohli, G. Kleywegt, E. Birney, D. Hassabis, and S. Velankar. 2022. AlphaFold Protein Structure Database: massively expanding the structural coverage of protein-sequence space with high-accuracy models. Nucleic Acids Res. 50:D439–D444.

Veatch, J.R., M.A. McMurray, Z.W. Nelson, and D.E. Gottschling. 2009. Mitochondrial dysfunction leads to nuclear genome instability via an iron-sulfur cluster defect. Cell. 137:1247–1258.

Vogtle, F.N., J.M. Burkhart, H. Gonczarowska-Jorge, C. Kucukkose, A.A. Taskin, D. Kopczynski, R. Ahrends, D. Mossmann, A. Sickmann, R.P. Zahedi, and C. Meisinger. 2017. Landscape of submitochondrial protein distribution. Nat Commun. 8:290.

Wurm, C.A., and S. Jakobs. 2006. Differential protein distributions define two sub-compartments of the mitochondrial inner membrane in yeast. FEBS Lett. 580:5628–5634.

Zhang, S., B. Reljic, C. Liang, B. Kerouanton, J.C. Francisco, J.H. Peh, C. Mary, N.S. Jagannathan, V. Olexiouk, C. Tang, G. Fidelito, S. Nama, R.K. Cheng, C.L. Wee, L.C. Wang, P. Duek Roggli, P. Sampath, L. Lane, E. Petretto, R.M. Sobota, S. Jesuthasan, L. Tucker-Kellogg, B. Reversade, G. Menschaert, L. Sun, D.A. Stroud, and L. Ho. 2020. Mitochondrial peptide BRAWNIN is essential for vertebrate respiratory complex III assembly. Nat Commun. 11:1312.

Zheng, X., and M. Xiang. 2022. Mitochondrion-located peptides and their pleiotropic physiological functions. FEBS J. 289:6919–6935.

